# Charge-Patterned Disordered Peptides Tune Intracellular Phase Separation in Bacteria

**DOI:** 10.1101/2023.08.01.551457

**Authors:** Jane Liao, Vivian Yeong, Allie C. Obermeyer

## Abstract

Subcellular phase separated compartments, known as biomolecular condensates, play an important role in the spatiotemporal organization of cells. To understand the sequence-determinants of phase separation in bacteria, we engineered protein-based condensates in *Escherichia coli* by utilizing electrostatic interactions as the main driving force. Minimal cationic disordered peptides were used to supercharge negative, neutral, and positive globular model proteins, enabling their phase separation with anionic biomacromolecules in the cell. The phase behavior was governed by the interaction strength between the cationic proteins and anionic biopolymers in addition to the protein concentration. The interaction strength primarily depended on the overall net charge of the protein, but the distribution of charge between the globular and disordered domains also had an impact. Notably, the protein charge distribution between domains could tune mesoscale attributes such as the size, number, and subcellular localization of condensates within *E. coli* cells. The length and charge density of the disordered peptides had significant effects on protein expression levels, ultimately influencing the formation of condensates. Taken together, charge-patterned disordered peptides provide a platform for understanding the molecular grammar underlying phase separation in bacteria.

**Highlights:**

- Minimal disordered cationic peptides of varying charge densities can promote protein phase separation in bacterial cells.
- Protein net charge and charge-patterning are distinct determinants of phase behavior.
- Protein charge distribution can be used to tune the size, number, position, and reversibility of condensates.
- Multiple proteins can be selectively recruited to synthetic condensates with disordered cationic peptides.

## Introduction

The compartmentalization of biomolecules is crucial for the spatiotemporal organization of cells. In addition to the wellknown membrane-bound organelles that serve diverse functions in eukaryotic cells, a host of naturally occurring membraneless intracellular compartments have been discovered and characterized in eukaryotes (1). Commonly referred to as biomolecular condensates, these intracellular compartments often concentrate proteins and nucleic acids and play important roles in biological processes spanning molecular to cellular length scales (2). New insights into the mechanism of their formation over the last few decades have shed light on numerous endogenous condensates in eukaryotes, including the nucleolus, Cajal bodies, nuclear speckles, paraspeckles, stress granules, and P granules (3–9). These phase-separated assemblies have been implicated in a variety of essential cellular processes ranging from regulating cell signaling to redirecting metabolic flux and adapting to stresses. Recent work suggests that condensates also serve an equally important role in the organization of the bacterial cytoplasm (10–15). However, despite significant progress towards understanding the formation, composition, material properties, and biological functions of condensates in eukaryotic cells, such insights are still largely lacking in bacterial cells.

The formation mechanism of bacterial condensates can be elucidated by borrowing concepts from eukaryotic counterparts, where weak multivalent interactions between constituent biomolecules are central to driving condensate assembly (16). Here, valency refers to the number of interaction sites, which can be surface regions on a folded protein, or amino acid motifs in an intrinsically disordered region or protein (IDR, IDP). A simple yet powerful framework for describing multivalent biomolecules is the stickers-and-spacers model from polymer physics, where stickers are defined as specific interaction sites that form physical crosslinks through noncovalent interactions such as hydrophobic, cation-π, aromatic, and electrostatic (17–19). Importantly, multivalent biomolecules above a threshold concentration, *c*_*perc*_, can undergo a networking transition known as bond percolation. Similarly, above a threshold concentration *c*_*sat*_, biomolecules can undergo a density transition whereby specific biomolecules de-mix into coexisting dilute and dense phases. Accordingly, phase separation aided bond percolation transitions have been proposed as a potential mechanism for condensate formation and dissolution in eukaryotic and bacterial cells (20).

Compositionally, condensate constituents are broadly comprised of scaffolds that drive phase separation and clients that are recruited through interactions with the scaffold biomolecules. Studies of both endogenous and synthetic eukaryotic condensates have primarily focused on IDRs as condensate scaffolds (21). In contrast to folded globular domains, the hallmark of IDRs is a high degree of conformation heterogeneity, which enables more flexible interaction modes. A subset of IDRs known as low-complexity domains show compositional bias toward a small set of amino acid stickers such as aromatic or charged residues (22). Although IDRs are ubiquitous in eukaryotic proteomes, comprising around 33% of the proteome, they are relatively scarce in bacteria and make up around 4% of bacterial proteomes (23). Despite this, IDRs have been identified as drivers of phase separation in prominent examples of endogenous bacterial condensates, including the disordered carboxyl-terminal (C-terminal) linker of FtsZ, a key bacterial cell division protein (24, 25), and the disordered C-terminal domain of *Caulobacter crescentus* RNase E, an essential endoribonuclease involved in RNA degradation (26, 27).

Given the importance of IDRs and biomolecular condensates to molecular functions and cellular processes in bacteria, several studies have engineered synthetic IDP-based systems, often with simplified repetitive sequence motifs, to uncover the sequence-determinants and molecular grammar of bacterial condensates. Examples in *Escherichia coli* (*E. coli*) include the recombinant overexpression of IDPs such as elastin-like polypeptides, spider silk, and resilin to form membraneless compartments (28, 29), the *de novo* engineering of condensates using artificial IDPs with different molecular weights and aromatic content (30), and the use of synthetic resilin-like polypeptides fused to functional domains to increase gene transcription by enriching plasmids and key components of the transcriptional machinery within condensates (31).

Although significant progress has been made in understanding the sequence heuristics of IDP driven phase separation in bacterial cells, most studies have predominantly focused on IDPs or IDRs while overlooking contributions from globular protein domains to the overall phase behavior. Previous work has shown that increasing the surface charge of globular proteins can enable phase separation with nucleic acids in bacteria via electrostatic interactions, even in the absence of a disordered domain (32). Considering that many endogenous condensate scaffolds consist of IDRs tethered to folded functional domains, we engineered modular proteins with a globular domain and a C-terminal charge-patterned disordered peptide domain. The use of minimal cationic peptides allowed us to preserve the relative and contextual importance of the globular domain in *de novo* condensates. We explored the interplay of contributions from both domains by systematically constructing a panel of proteins from three globular green fluorescent proteins (GFP), derived from superfolder GFP, with different surface charges and two cationic disordered peptide motifs with varied charge-patterning.

Specifically, we explored the charge, charge-patterning, and concentration dependencies of intracellular phase separation by overexpressing this panel of engineered proteins in *E. coli* to form *de novo* condensates. We show that the formation of electrostatically driven condensates depends primarily on the net charge and *in vivo* concentration of the protein scaffold. Important sequence-specific determinants such as amino acid composition, charge-patterning, and peptide length altered the interaction strength and influenced protein expression levels. Interestingly, fluorescence imaging revealed that sequence-encoded features could tune mesoscale attributes such as the size, number, and subcellular localization of condensates within *E. coli* cells, alluding to intricate structure-form-function relationships that underly endogenous condensates. Taken together, the use of short disordered peptides is a versatile approach to expand our fundamental understanding of the sequence parameters governing biomolecular condensates in bacteria.

## Results

### Design of disordered cationic peptides with varying charge densities

Along with an abundance of anionic biomacromolecules such as RNA and DNA, the *E. coli* proteome is also comprised of negatively charged proteins (Figure 1A). This allows us to leverage electrostatic attraction and entropic gains from counterion release to drive the partitioning of cationic proteins of interest in complex coacervates. It was previously demonstrated that the negative globular GFP could be supercharged with a C-terminal cationic peptide to form phase separated intracellular condensates without the need to introduce additional anionic partners (33). Recent mounting evidence has also revealed the importance of the arrangement of residues in the primary sequence in modulating the strength of the weak, multivalent interactions responsible for intracellular phase separation (33–36). To explore the role of charge-patterning in the complex coacervation of proteins in cells, we designed two cationic peptides with differing charge densities (Figure 1B and Figure S1). The low charge density peptide (L_n_) has the amino acid sequence of GRRGKKSRK, with neutral glycine and serine residues interspaced between every 2 cationic residues, while the high charge density peptide (H_n_) has the sequence KRRRKK and only consists of cationic residues. Repeats of the base peptide sequences were also studied to investigate the effects of increasing peptide length, which alters both the net protein charge and the balance of charge between the disordered peptide domain and the globular protein domain. Here, we also probed the relative importance of electrostatic interactions from both folded and disordered domains by appending the low and high charge density cationic peptides to engineered GFP variants with anionic (GFP(−6)), neutral (GFP(0)), or cationic (GFP(+6)) net charge. In addition to the presence of phase separation within a cell population, we aimed to elucidate the parameters governing the number, size, and protein partitioning of engineered condensates (Figure 1C). As a more conservative estimate of protein partitioning, we calculated the average fluorescence intensity of condensates relative to the cytoplasm rather than comparing the maximum intensity within condensates to the cytoplasm. We hypothesized that both the condensate size and strength of protein partitioning would depend on the net charge and intracellular protein concentration. We also postulated that proteins with equivalent net charge could have different propensities for homotypic and heterotypic interactions, based on whether the charge resides primarily on the folded or disordered domain.

**Figure 1.**
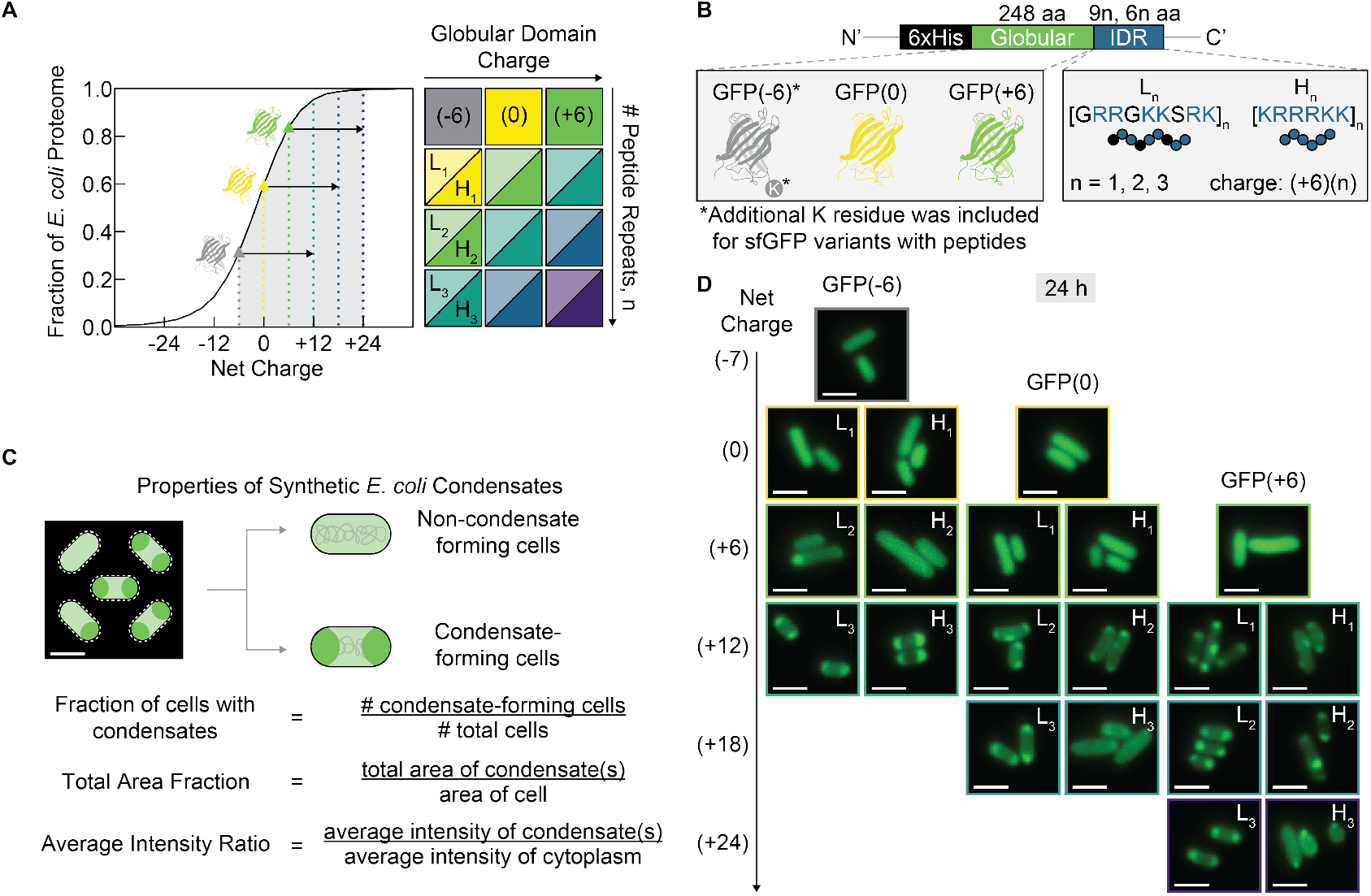
Design of disordered cationic peptides with varying charge density to create synthetic biomolecular condensates in *E. coli* (A) Cumulative distribution of proteins in the *E. coli* proteome (UP000002032) by expected charge. Triangular markers indicate the predicted charge of engineered isotropic proteins and arrows indicate the net charge increase from disordered cationic peptides. (B) Schematic for the design of globular and C-terminal charge-patterned disordered peptide domains. (C) Cell population, cell, and condensate properties assessed from fluorescence microscopy images. (D) Representative microscopy images of cells at 24 h post-induction shown according to overall net charge. GFP(−6)-L_2_ and all protein variants with net charge ≥ +12 form foci within the cell. Scale bars are 2 μm.

### The role of charge-patterning in promoting bacterial condensate formation

In cases when the protein interaction strength is sufficient for associative heterotypic phase separation, condensates should form if cellular levels of the protein are greater than the saturation concentration, *c*_*sat*_. Given the hypothesized importance of the intracellular protein concentration in enabling phase separation, the fluorescence intensity normalized by the cell density (FI/OD_600_) was used as a proxy to estimate the mean cell protein concentration. At 24 h post-induction, both the OD_600_ and the FI/OD_600_ were found to decrease with increasing peptide charge density and length, suggesting that long charge-dense disordered peptides have a significant effect on the host cell growth and protein expression levels (Figure 2A, Figure S2A, and S2B).

**Figure 2.**
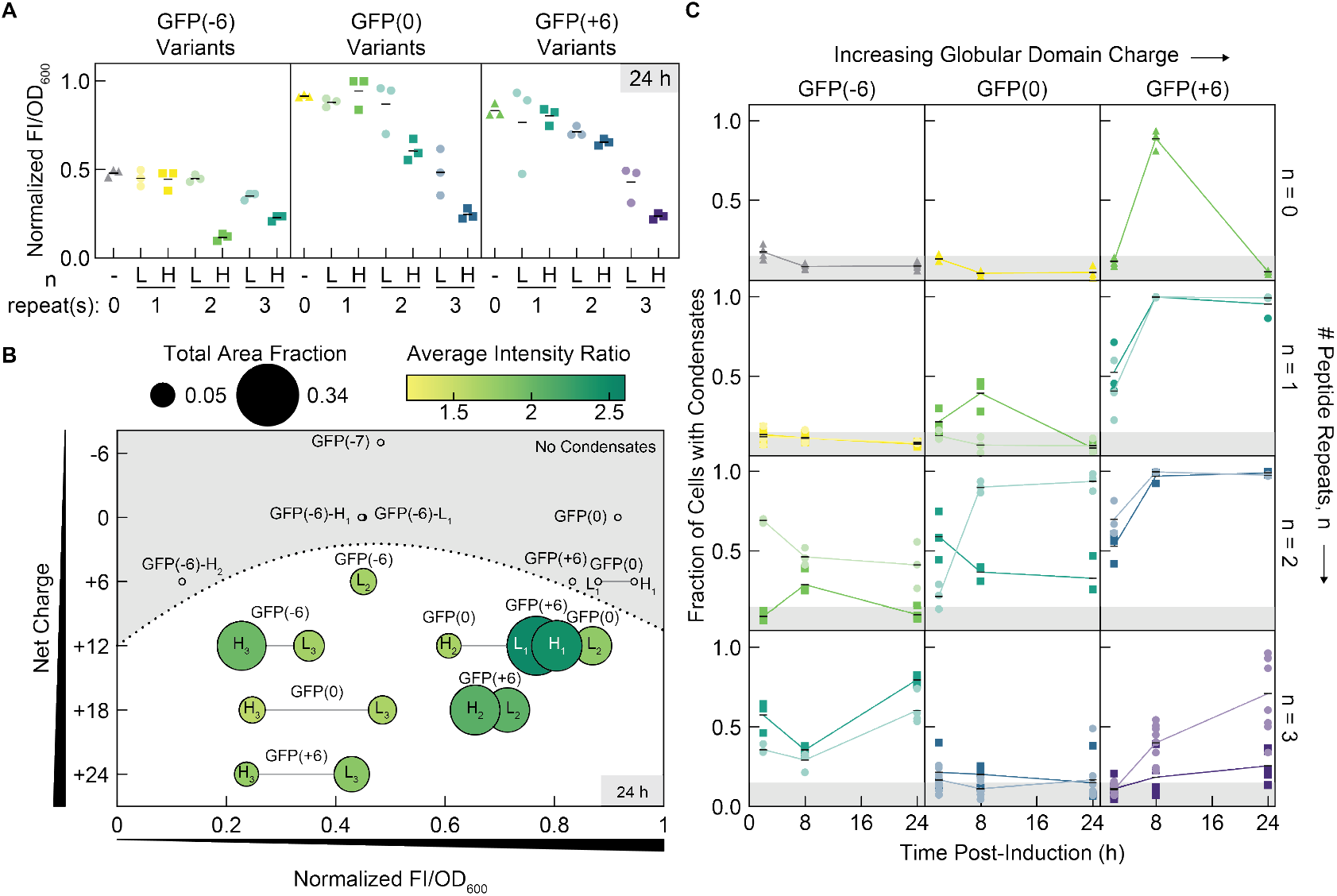
Overall net charge, charge distribution, peptide charge-patterning, and protein concentration determine the phase separation behavior of GFP variants. (A) The ratio of fluorescence intensity (FI) to cell density (OD_600_) was used as a proxy for intracellular GFP concentration. At 24 h post-induction, the normalized FI/OD_600_ decreases with both peptide charge density and peptide length. Data was normalized to GFP(0)-H_1_ at 24 h as the maximum. Three biological replicates and the respective means are shown. Triangles, circles, and squares indicate isotropic variants, low charge density peptide variants, and high charge density peptide variants, respectively. (B) At 24 h post-induction, the total area fraction and median intensity ratio of condensates depend on the net charge and expression of the protein. Variants with high net charge and FI/OD_600_ tend to form larger and brighter condensates. Marker size and shading indicates the median total area fraction and median average intensity ratio in cells with condensates. Grey lines show the pairwise comparisons between different charge density peptides on the same globular domain. Grey circles mark the FI/OD_600_ for variants that do not form condensates at 24 h post-induction. (C) Variants with a net charge of -7 or 0 do not form condensates at 2, 8, or 24 h post-induction. Some variants with a net charge of +6 exhibit reversible condensate formation over time. The mean and a minimum of three biological replicates for each variant are shown. Four biological replicates are shown for sfGFP, GFP(−6)-H_2_, GFP(−6)-L_3_, GFP(0)-H_3_, and GFP(+6); five biological replicates are shown for GFP(+6)-H_3_ ; and eight biological replicates are shown for GFP(0)-L_3_ and GFP(+6)-L_3_. At least 83 cells were analyzed per biological replicate. Triangles, circles, and squares indicate isotropic variants, low charge density peptide variants, and high charge density peptide variants, respectively.

To investigate the effects of charge-patterning on intracellular phase separation facilitated by these short, disordered peptides, each GFP variant was expressed in *E. coli* cells and monitored by fluorescence microscopy for the presence of bacterial condensates (Figure 1D). Consistent with previous observations, the net charge was found to be a main governing parameter for condensate formation, and all variants with net charge ≥ +12 showed observable condensates at 24 h post-induction (33). Interestingly, GFP(−6)-L_2_ also demonstrated intracellular phase separation at 24 h post-induction despite having a net charge of +6. The existence of a critical net charge for phase separation was also evident in *in vitro* turbidity assays using purified protein and total RNA from torula yeast (Figure S2F and S2G). Analysis of the size of and protein partitioning in condensates according to the net charge and intracellular protein concentration revealed only minor differences with respect to the peptide sequence at 24 h post-induction for variants with the GFP(+6) domain (Figure 2B). This suggests that small sequence modifications on the disordered peptide domain have minor effects at long expression times if the globular domain also facilitates phase separation.

Although differences owing to the charge density of the peptide were largely overshadowed when appended to a positive globular domain, the charge-patterning of the peptide had noticeable effects when tethered to negative or neutral globular domains. Notably, there were differences in the size and fluorescence intensities between GFP(−6)-L_3_ and GFP(−6)-H_3_ condensates, as well as between GFP(0)-L_2_ and GFP(0)-H_2_ condensates. As neither the GFP(−7) nor GFP(0) globular domains phase separated without disordered cationic peptides, the peptide was likely the main driver of phase separation for these proteins by facilitating multivalent interactions with anionic macromolecules. Consequently, the primary sequence of the peptide is particularly important for negative or neutral globular domains. In contrast, the peptide charge-patterning is of less importance when both the globular and peptide domains have the potential for complex coacervation, as with the GFP(+6) variants fused to cationic peptides. Irrespective of the peptide charge density, GFP(+6) protein variants formed larger and brighter condensates at 24 h post-induction when compared to GFP(−6) or GFP(0) variants with equivalent net charge.

Quantitative image analysis further revealed that the fraction of condensate-forming and non-condensate-forming cells changes in the late-log, early-stationary, and late-stationary phases of cell growth at 2, 8, and 24 h post-induction, respectively (Figure 2C). Variants with high charge density peptides at the net charge threshold of +6 formed reversible condensates, whereby the fraction of cells with condensates increased from 2 to 8 h and decreased from 8 to 24 h postinduction. Condensate reversibility was previously demonstrated with the isotropic GFP(+6) variant, and these findings provide additional evidence that anisotropic proteins with disordered peptides can also undergo similar reversible condensate formation. In general, longer peptide lengths appeared to enable a moderate fraction of the cell population to form condensates even as early as 2 h post-induction, as with GFP(−6)-L_2_, GFP(−6)-L_3_, GFP(−6)-H_3_, GFP(0)-H_2_, GFP(+6)-L_2_, and GFP(+6)-H_2_. We hypothesized that this may be due to an increase in the protein charge patchiness and the radii over which the charge extends. For increasingly longer peptide lengths, the reduction of protein expression likely counteracted the increase in interaction affinity. This is shown for GFP(0) and GFP(+6) with 3 repeated motifs, which have lower propensity to form condensates compared to respective variants with 2 repeated motifs. Among the complete charge matrix of variants that were able to form condensates, we chose to examine variants with net charges of +6 and +12 in more detail to understand the effects of charge distribution and peptide charge density at and above the charge threshold for intracellular phase separation in *E. coli*.

### Reversible condensate formation at the charge threshold is affected by charge isotropy

It was previously observed that at a net charge of +6, both isotropic variants and variants with a cationic disordered peptide had the propensity to undergo reversible phase separation. This behavior was shown to correlate with the growth phases, with condensates forming during the late-log phase and disassembling during the stationary phase. Here, fluorescence microscopy revealed similar condensate reversibility among the variants with high charge density peptide domains (Figure 3A and B). The fraction of cells with condensates increased throughout the late log-phase until 8 or 9 h post-induction (Figure 3E). The maximum fraction of cells with condensates differed among the variants according to the protein charge distribution. A very low fraction of cells expressing GFP(−6)-H_2_ formed condensates by 8 h post-induction, while around half of all cells expressing GFP(0)-H_1_ formed condensates by 9 h, and nearly all cells expressing isotropic GFP(+6) formed condensates by 8 h.

**Figure 3.**
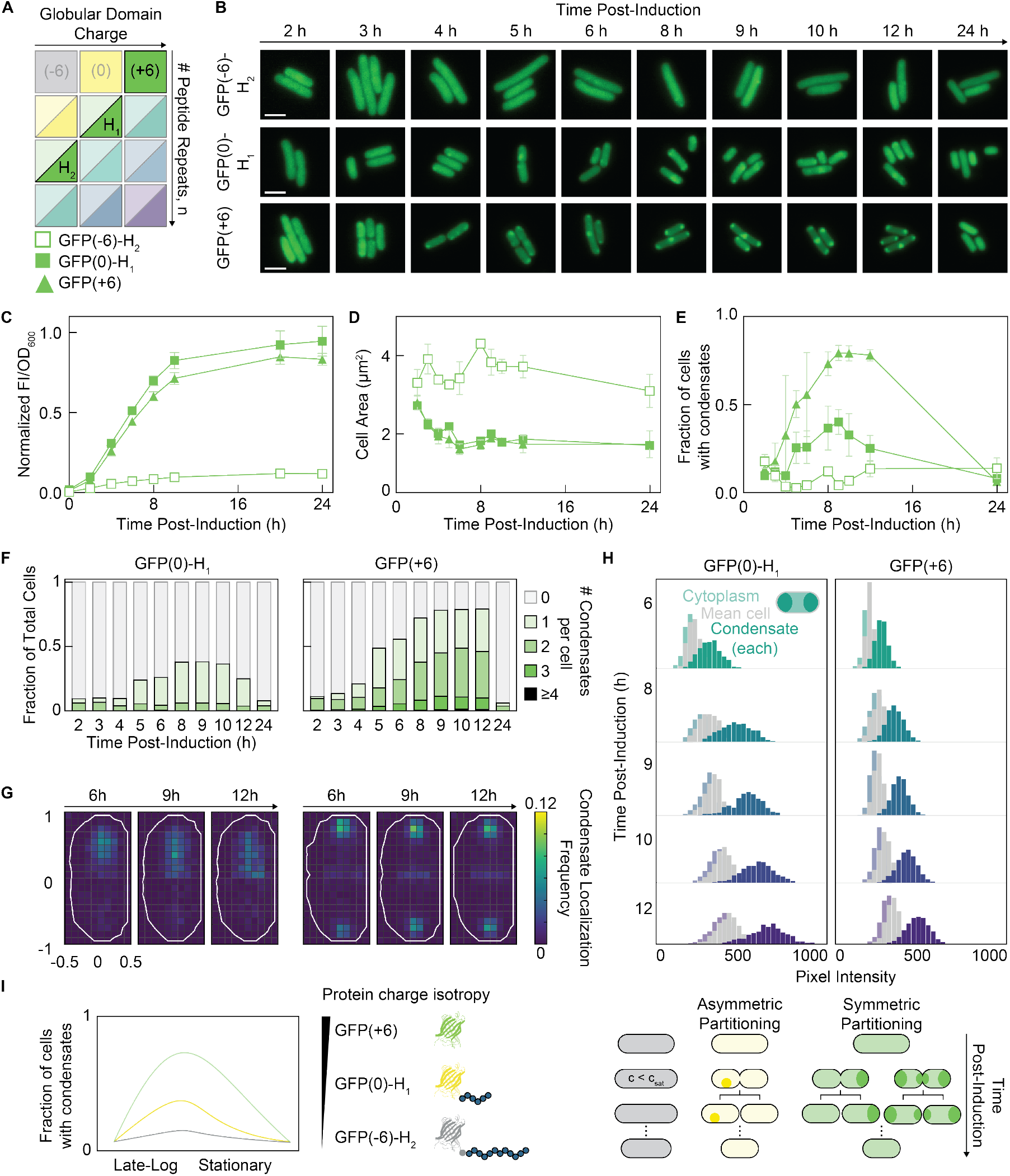
Variants with net charge of +6 are at the net charge threshold for condensate formation and exhibit condensate reversibility over time. The spatiotemporal properties of condensates vary with the charge distribution between the globular protein and disordered peptide domains. (A) Schematic of the experimental charge matrix with the +6 net charge high charge density peptide variants highlighted. (B) Representative time-course microscopy images of cells from 2 to 24 h post-induction for GFP(−6)-H_2_, GFP(0)-H_1_, and GFP(+6). All three variants show the formation and dissolution of condensates over time. Scale bars are 2 μm. (C) Normalized FI/OD_600_ over time for GFP(−6)-H_2_, GFP(0)-H_1_, and GFP(+6). Data points and error bars indicate the mean and standard deviation of the three biological replicates. Lines connecting the means of three biological replicates are shown. (D) Cells expressing GFP(−6)-H_2_ are approximately twice as large as cells expressing GFP(0)-H_1_ or GFP(+6). The mean and standard deviation of three biological replicates are shown. (E) The peak fraction of cells with condensates is higher and the duration of condensate formation increases for more isotropic charge distributions. At least 134 cells were analyzed per replicate and timepoint. The mean and standard deviation of three biological replicates are shown. (F) Among condensate-forming cells, a majority of GFP(0)-H_1_ cells form 1 condensate per cell while a majority of GFP(+6) cells form ≥ 2 condensates per cell. Data from all biological replicates were consolidated. (G) As shown by binned, normalized, and aligned 1-pixel local maxima heat maps of all condensate-forming cells, GFP(0)-H_1_ condensates (left) do not have a strong spatial localization preference, while GFP(+6) condensates (right) preferentially localize at the cell poles in shorter cells and at the cell poles and midcell in longer cells. Outlines depict an approximate cell contour. (H) The pixel intensity distribution of GFP(0)-H_1_ condensates show a larger spread over time when compared to GFP(+6) condensates, suggesting greater cell-to-cell variability of condensates when the short peptide domain is the primary driver of phase separation. Data is shown for the condensate-forming subpopulation of consolidated biological replicates with a histogram bin width of 30. (I) Volume and shape changes during the cell cycle may influence the formation and dissolution of condensates by concentrating or diluting the protein within the cell. The initial cell size increase of GFP(−6)-H_2_ may hinder phase separation by decreasing the protein concentration below the saturation concentration. The number and localization of condensates determine whether condensates will be asymmetrically or symmetrically inherited from to mother to daughter cell.

The lower prevalence of condensates among cells expressing GFP(−6)-H_2_ may be attributed to significantly lower protein expression compared to GFP(0)-H_1_ and GFP(+6) (Figure 3C and Figure S3). Interestingly, cells expressing GFP(−6)-H_2_ initially increased in cell area during the late-log phase and were approximately twice as long as cells with GFP(0)-H_1_ or GFP(+6) in the stationary phase (Figure 3D). As the late-log phase was shown to be a critical period for initial condensate formation, we hypothesized that the dilution of protein via cell elongation could have further prevented the formation of GFP(−6)-H_2_ condensates. To examine this further, we characterized condensate properties such as number, localization, and partitioning for GFP(0)-H_1_ and GFP(+6). These two variants had very similar protein expression levels and cell area over time, enabling a direct comparison with fewer confounding variables such as intracellular protein concentration (Figure 3C and D). Interestingly, among condensate-forming cells, GFP(0)-H_1_ preferentially formed one condensate per cell while GFP(+6) had a tendency to form two or more condensates per cell (Figure 3F). Given similar intracellular protein concentrations between these two variants, the greater number of condensates per cell for GFP(+6) may be attributed to more potential nucleation sites and a wider phase separation window. It is known that a higher connectivity between multivalent molecules is key to network formation and phase separation (34, 35). A positive isotropic globular domain potentially provides more discrete nodes to initiate and support system-spanning networks when compared to a short, localized, charge-dense peptide domain. In addition to differences in the number of condensates per cell, GFP(0)-H_1_ condensates also did not exhibit preferential subcellular localization within a given cell compared to GFP(+6) which strongly localized to both cell poles. In longer cells, GFP(+6) also had a strong preference for a third foci localization at the midcell (Figure 3G).

The number and localization of condensates can influence the inheritance of condensates from mother to daughter cell, which dictates the persistence of condensate-forming cells in a population. In prior work, the selective spatial localization of biomolecules has been shown to modulate the differential cell fate of daughter cells and population-level cellular behaviors (31, 36–38). For example, engineering the asymmetric segregation of plasmid DNA to a single position within *E. coli* can lead to cells with distinct differentiated states (36). Similarly, we observed that single polarly-localized condensates were typically asymmetrically inherited during cell division whereby only one of the daughter cells retained the condensate (Figure 3I). This limited the maximum fraction of condensate-containing cells in the population to roughly half of the cell population, as with GFP(0)-H_1_ at 8 h post-induction. Conversely, cells expressing GFP(+6) preferentially formed condensates at both cell poles, resulting in symmetric inheritance. It was noted that dividing cells expressing GFP(+6) also had a tendency to form a third condensate at the midcell, in addition to condensates at the poles. In this case, both daughter cells could inherit two condensates. Regardless, GFP(+6) condensates were both more prevalent and persistent in the cell population, suggesting that simply altering the distribution of net charge of the protein scaffold could have broader population-level consequences.

In addition to differences in the number and localization of condensates, the degree of protein partitioning between the condensed phase and the surrounding cytoplasm was also investigated for the GFP(0)-H_1_ and GFP(+6) variants. In agreement with the FI/OD_600_ data, the mean cell fluorescence, as calculated by the total cell fluorescence normalized by the cell area, increased slightly from 6 h to 12 h post-induction (Figure 3H). The distributions of mean cytoplasm intensity and mean condensate intensity also increased as a function of this. Although the average intensity ratio, comparing condensate to cytoplasm intensity, did not change significantly from 6 h to 12 h post-induction, ∼ 1.6-1.7 for GFP(0)-H_1_ and ∼ 1.5-1.6 for GFP(+6), greater cell-to-cell variability was seen in the broadening condensate intensity distribution of GFP(0)-H_1_. This suggests that the earlier disassembly of GFP(0)-H_1_ condensates is a result of exiting a narrower phase separation window.

Overall, the results suggest that even at similar protein expression levels, the phase separation behavior of a protein depends on whether the charge is isotropically distributed on the globular domain or resides entirely on a highly charge-dense peptide domain. It is also important to note that condensates must be considered in the context of the cell as a changing surrounding environment. In a typical batch culture, both mRNAs and select proteins are down-regulated in the nutrient-limited stationary phase as ribosomes become inactive and undergo ribosome hibernation (39, 40). It is well-known that many other morphological and physiological changes accompany the transition from the log-phase to the stationary phase of cell growth. These include cell volume decrease, cell shape change, nucleoid compaction, cell wall thickening, and cytoplasm composition change (41, 42). On a single-cell level, proteins may also be diluted by cell elongation or concentrated by cell division as cells undergo rounds of cell division.

### Protein scaffolds with higher net charge maintain condensates at later timepoints

In addition to protein scaffolds with a net charge of +6, protein variants with a higher net charge of +12 and longer disordered peptide domains were also studied to elucidate the effects of charge and charge distribution on condensate properties (Figure 4A). Microscopy images revealed that condensates were present at 2, 8, and 24 h post-induction (Figure 4B). In contrast with most +6 variants that did not have condensates at 24 h postinduction, all +12 variants still had a moderate to high fraction of cells with condensates at longer timepoints, which may be attributed to greater overall net charge or longer peptide lengths, resulting in a broader two-phase region.

**Figure 4.**
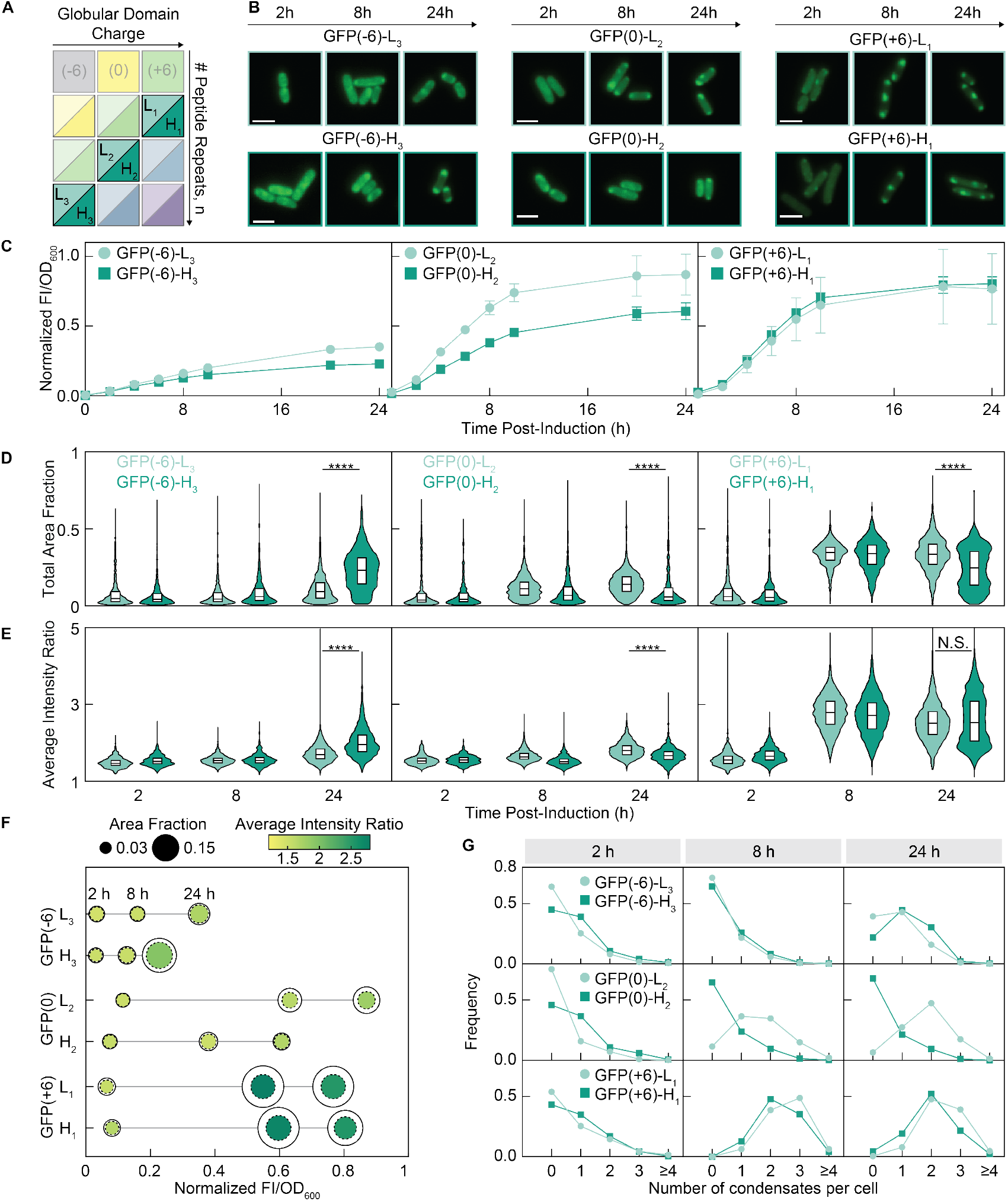
For variants with a net charge of +12, the peptide charge density and charge distribution between domains affects interaction strength and protein expression, and consequently the condensate area, intensity, and number of condensates per cell. (A) Schematic of the experimental charge matrix with the +12 net charge variants highlighted. (B) Representative microscopy images of cells at 2, 8, and 24 h post-induction for each variant. All variants with net charge ≥ +12 still have cells with foci at 24 h post-induction. Scale bars are 2 μm. (C) For negative GFP(−6) and neutral GFP(0) globular domains, appending a high charge density polypeptide results in lower normalized FI/OD_600_. Data was normalized to GFP(0)-L_2_ at 24 h post-induction as the maximum. Lines connecting the means of three biological replicates (data points) are shown. (D) The total area fraction of condensates increases over time. The total area fractions of condensates formed with proteins that have cationic charge on both the globular and disordered peptide domains are larger than for protein variants that only have cationic charge on the peptide domain. (E) The intensity ratio of condensates was used as a proxy for the degree of protein partitioning. On a negative globular domain, a high charge density peptide results in more protein partitioning in the condensate. The converse is true on a neutral domain, where a low charge density peptide led to more protein partitioning. On a positive globular domain, the partitioning is less affected by the charge density of the disordered peptide. GFP(+6) variants show higher protein partitioning. For (D) and (E), data from condensate-forming cells from all biological replicates were consolidated and each condensate was analyzed separately. At least 725 condensates were analyzed for each distribution. L peptide variants are light teal and H peptide variants are dark teal. Differences between variants at 24 h post-induction were evaluated by the Mann-Whitney test, with **** indicating p < 0.0001. (F) Summary data of the median area fractions and median intensity ratios of condensates as a function of the normalized FI/OD_600_ at 2, 8, and 24 h post-induction. The lower FI/OD_600_ of GFP(−6) variants may explain the smaller total area fraction and lower intensity ratios of the condensates at 8 h post-induction. Although the FI/OD_600_ of GFP(0) variants are similar to GFP(+6) variants, a neutral globular domain may have lower interaction strength compared to a positive globular domain, resulting in smaller and less bright condensates overall. Markers for each variant are in the order of time post-induction (2, 8, 24 h). Dashed circles indicate the median area fraction of individual condensates, and the solid circles indicate the median total area fraction of all condensates in a given cell. Although the total area fraction increases over time, the area fraction of individual condensates appear to have an upper bound. (G) The FI/OD_600_ may affect the number of condensates per cell. GFP(−6) variants have lower FI/OD_600_ at 24 h post-induction and the majority of condensate-forming cells have 1 condensate. GFP(0)-L_2_ has slightly higher FI/OD_600_ at 24 h post-induction and preferentially contains 2 condensates per cell, while GFP(0)-H_2_ has lower FI/OD_600_ and a majority do not form condensates. GFP(+6) variants have sufficient interaction strength and FI/OD_600_ to preferentially form 2 or 3 condensates per cell. Data from all biological replicates were consolidated.

The +12 variants could be categorized by the approximate intracellular protein concentration (Figure 4C and Figure S4). Specifically, GFP(−6)-L_3_, GFP(−6)-H_3_, and GFP(0)-H_2_ all had relatively lower protein expression levels. In the case of the GFP(−6)-L_3_/H_3_ variants, this was likely due to difficulty in expressing highly anisotropic proteins with long stretches of cationic residues. In the case of GFP(0)-H_2_, low expression may be due to the moderately long but highly charge dense H_2_ domain, as similar reduction in expression was seen previously with GFP(−6)-H_2_ (Figure 3C). The remaining variants, GFP(0)-L_2_, GFP(+6)-L_1_, and GFP(+6)-H_1_ did not exhibit significant reduction in protein expression when compared to the respective unmodified globular domains at 2, 8, or 24 h post-induction (Figure S2A).

Several condensate properties were consistent with the FI/OD_600_ or intracellular protein concentration. Variants with a higher FI/OD_600_ had condensates at 24 h post-induction that altogether occupied a greater fraction of the cell (Figure 4D). Interestingly, although the total area fraction of condensates increased substantially over time for variants with higher FI/OD_600_, the area fraction of individual condensates did not increase as significantly (Figure 4F). Instead, the increase in total area fraction occupied by condensates was a result of more condensates being formed per cell (Figure 4G). Using the total area fraction as a proxy for the total volume fraction of condensates, this suggests that although there is a driving force for increasing the total volume fraction of condensates, steric occlusion or other effects may prevent the formation of larger individual condensates or the fusion of multiple condensates.

The average partitioning of protein in condensates generally increased over time for all variants (Figure 4E). The effects of the peptide charge density were most noticeable on the negative GFP globular domain, with the high charge density H_3_ peptide leading to more protein partitioning in the condensed phase. The converse was true on a neutral domain, where a low charge density peptide led to more protein partitioning. On a positive globular domain, the average intensity ratios were less impacted by the charge density of the peptide. This supports the idea of a hierarchical relationship among the different sequence parameters, where the overall net charge and globular domain charge are important when the peptide length is short, while the peptide charge density is more relevant when the peptide is adequately long and serves as the primary interaction domain.

### Modular disordered cationic peptides sequester multiple proteins in engineered condensates

In addition to encapsulating a single protein of interest in phase separated condensates, the modular nature of the cationic peptides can be leveraged to colocalize multiple proteins within the same condensed phase. This allowed us to explore the sequence-determinants of multi-protein condensates that are more representative of endogenous systems. Here we demonstrate that multi-protein condensates consisting of GFP and another globular fluorescent protein, mCherry, can be engineered in *E. coli* cells with the low and high charge density peptide domains. Specifically, we co-expressed GFP(− 6)-H_2_ and mCherry(−6)-L_n_, where n represents the number of peptide motif repeats as before (Figure 5A). As a control, we first expressed mCherry(−6)-L_n_ alone, and confirmed that like GFP(−6)-L_n_, a net charge greater than +6 was required for mCherry condensate formation (Figure S5E, Figure 2C). Minor differences in the phase separation behavior between GFP(−6)-L_2_/L_3_ and mCherry(−6)-L_2_/L_3_ may be attributed to differences in protein expression or protein maturation times. We hypothesized that the characterized phase separation behavior of GFP and mCherry would largely remain the same even if they were present in multi-protein assemblies (Figure 5B).

**Figure 5.**
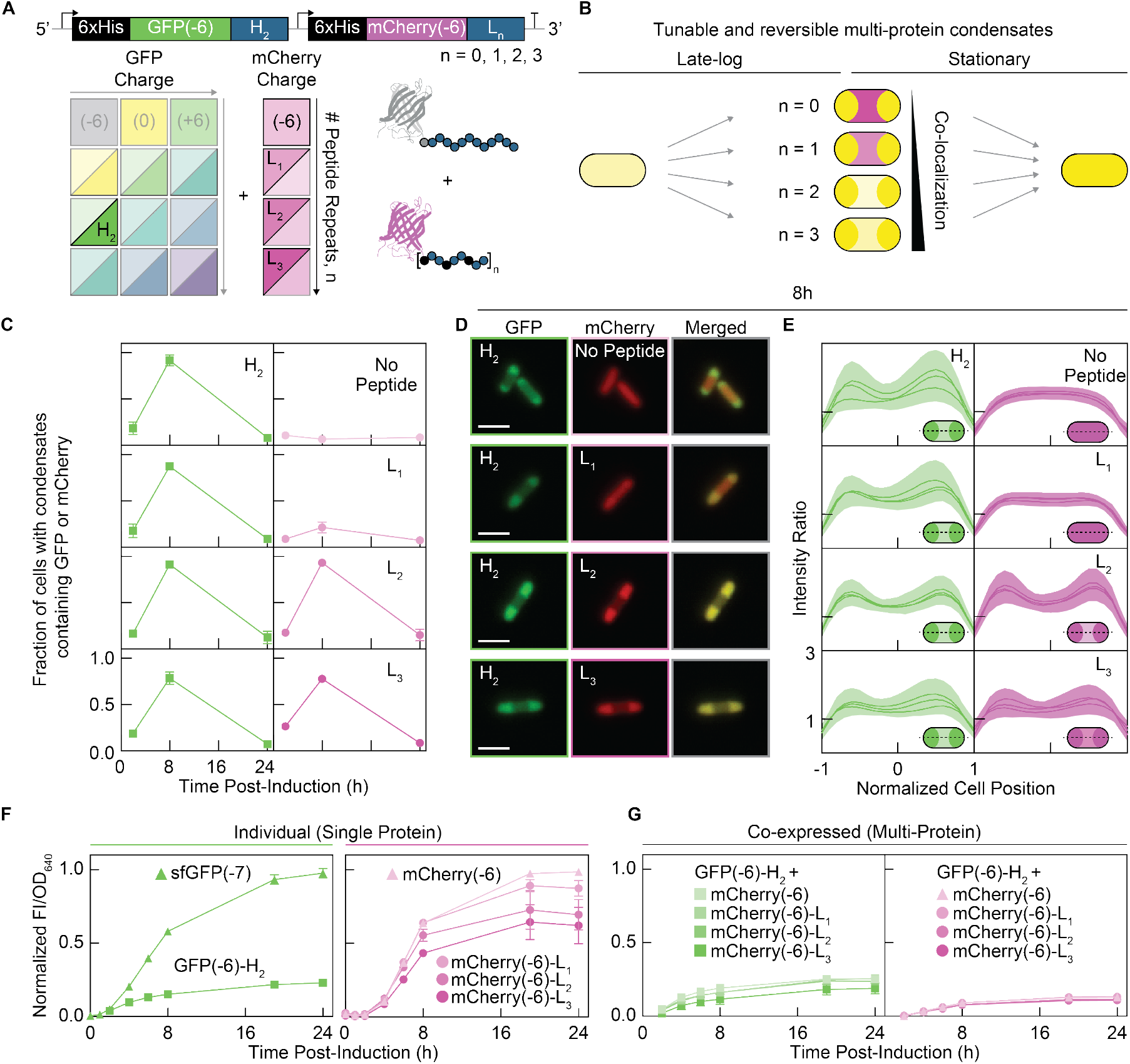
Modular disordered cationic peptides can be used to engineer reversible multi-protein condensates with GFP and mCherry. The degree of mCherry partitioning can be tuned with the length of the peptide. (A) GFP(−6)-H_2_ and mCherry-L_n_ were co-expressed from a single transcript by separate T7 promoters and ribosome binding sites. Experimental charge matrix shows the peptides used for each protein. (B) Schematic of the reversibility and tunability of multi-protein GFP and mCherry condensates. (C)The fraction of cells with condensates containing GFP(−6)-H_2_ is similar despite different co-expression partners, suggesting GFP(−6)-H_2_ acts as a scaffold. The fraction of cells with condensates containing mCherry-L_n_ is tunable and increases with the number of peptide motif repeats, n, at 8 h post-induction. Microscopy images from each fluorescence channel were analyzed independently. The fraction of cells with condensates from each channel were calculated separately. Mean and standard deviation of three biological replicates are shown. (D) Representative microscopy images at 8 h post-induction when nearly all cells form condensates with at least one protein. GFP(−6)-H_2_ and mCherry-L_n_ show stronger colocalization with increasing cationic peptide length on mCherry. Scale bar is 2 μm. (E) The intensity ratio along the medial axis shows preferential localization of GFP(−6)-H_2_ at the cell poles. mCherry-L_n_ colocalizes with GFP(−6)-H_2_ when it is charge equivalent or has slightly higher charge than GFP(−6)-H_2_. The intensity ratios along the medial axis were normalized to cell-position and averaged. For cells containing one condensate, the medial profile was aligned with the condensate on the positive end of the medial axis. Lines and shading indicate the mean and standard deviation of three biological replicates. Microscopy images from each fluorescence channel were analyzed independently. (F) FI/OD_640_ for cells expressing a single protein show that the high charge density H_2_ peptide significantly reduced the protein expression of GFP(−6)-H_2_ compared to sfGFP. Increasing repeats of the low charge density peptide sequence moderately reduced the expression of mCherry-L_n_ when compared to mCherry. Data for GFP and mCherry variants were normalized to sfGFP(−7) at 24 h and mCherry(−6) at 24 h as the maximum. The mean and standard deviation for three biological replicates are shown. GFP and mCherry fluorescence were measured with excitation/emission of 470/510 nm and 570/610 nm, respectively. OD was measured at 640 nm. (G) When co-expressed, both GFP(−6)-H_2_ and mCherry-L_n_ show similar expression levels to GFP(−6)-H_2_ when expressed alone. Data for GFP and mCherry variants were normalized to sfGFP(−7) at 24 h and mCherry(−6) at 24 h as the maximum. The mean and standard deviation for three biological replicates are shown.

As such, we expected GFP(−6)-H_2_ to still undergo reversible phase separation over time when co-expressed (Figure 2C). Consistent with our expectations, GFP(−6)-H_2_ formed reversible condensates regardless of the net charge of mCherry, although a much higher fraction of cells contained GFP(−6)- H_2_ condensates at 8 h post-induction (Figure 2C and Figure 5C). Given that RNA length has been shown to modulate condensates (43, 44), a potential explanation may be that the longer mRNA transcripts can facilitate more associative interactions with GFP(−6)-H_2_ to promote phase separation. In the case of co-expression with mCherry(−6)-L_2_ or mCherry(− 6)-L_3_, the synergistic effects of expressing two proteins that are independently capable of phase separation may also help promote GFP(−6)-H_2_ condensation.

Similarly, mCherry(−6), mCherry(−6)-L_1_, and mCherry(−6)- L_2_ also behaved in a predictable manner when co-expressed with GFP(−6)-H_2_ (Figure S5E and Figure 5D). In the case of mCherry(−6) and mCherry(−6)-L_1_, neither protein had sufficient charge to phase separate when expressed alone or when co-expressed with GFP(−6)-H_2_, as shown by the average intensity ratio along the medial-axis of the cell (Figure 5E). Co-expression of charge-equivalent mCherry(−6)- L_2_ and GFP(−6)-H_2_ revealed that mCherry(−6)-L_2_ behaved nearly identically to GFP(−6)-H_2_ in terms of condensate prevalence among the cell population, localization to the cell poles, and formation and disassembly over time. Interestingly, co-expression with GFP(−6)-H_2_ appeared to slightly modify the phase separation behavior of mCherry(−6)-L_3_. Although when expressed alone, mCherry(−6)-L_3_ condensates were present after 24 h post-induction, mCherry(−6)-L_3_ condensates seemed to disassemble earlier along with GFP(−6)- H_2_ condensates when co-expressed. Taken together, both the invariable phase behavior of GFP(−6)-H_2_ despite different coexpression partners and the modulating effect it has on the phase behavior of mCherry(−6)-L_3_ suggests that it acts as a scaffold, potentially due to the earlier upstream expression of GFP(−6)-H_2_, the higher charge density of the H_2_ peptide domain, or a slightly higher expression level (Figure 5G).

## Discussion

Here, we explored the effects of charge, charge-patterning, and concentration on electrostatically driven intracellular phase separation in *E. coli* using engineered protein scaffolds consisting of a globular domain and a charge-patterned disordered peptide domain. The overall net charge and intracellular protein concentration were shown to be important parameters governing the heterotypic phase separation of cationic protein scaffolds with anionic biomolecules in the cell. Proteins with negative or neutral net charge did not phase separate *in vivo*, while most proteins with a net charge ≥ +6 had sufficient valence to form condensates. The patterning of charged residues in the disordered peptide domain had a significant effect on protein expression, and a more charge-dense motif noticeably reduced the intracellular protein concentration. At the phase separation net charge threshold of +6, variants had the propensity to undergo reversible phase separation, whereby condensates formed during the late-log phase of cell growth and disassembled in the stationary phase of cell growth. At a fixed protein net charge of +6, altering the distribution of charge between the folded globular domain and the disordered cationic peptide domain led to differences in mesoscale features such as the size, number, and subcellular localization of condensates.

Protein variants with a higher net charge of +12 formed condensates that could be maintained at longer timepoints, corresponding to phase separation over a broader range of protein concentrations. When the disordered peptide domain was the sole contributor of the net cationic charge, as with GFP(−6)-L_3_ and GFP(−6)-H_3_, the peptide charge density had a noticeable effect on the size and partitioning of proteins in the condensates. Specifically, the higher charge density KRRRKK peptide motif led to slightly larger condensates and higher protein partitioning than the GRRGKKSRK peptide motif. Taken together, this suggests that when the disordered domain is the primary driver of phase separation, highly charge-dense sequences may increase the affinity of the protein towards anionic biomolecules to facilitate phase separation. However, this is balanced by a counteracting effect whereby highly charge-dense sequences negatively impact cell physiology and protein expression levels and consequently impede phase separation.

Although understanding the design parameters governing the intracellular phase separation of a single protein is invaluable, endogenous biomolecular condensates in bacteria are multicomponent systems consisting of numerous proteins and nucleic acids. As such, we also used the modular nature of cationic peptides to elucidate the sequence-determinants of the phase behavior and partitioning of two globular proteins, GFP and mCherry, in multi-protein systems. We specifically studied cases in which the cationic peptide was the main driver of phase-separation by co-expressing GFP(−6)-H_2_ with mCherry(−6)-L_n_. We varied the valence, or net charge, of mCherry by changing the length of the attached disordered peptide. Consistent with our previous observations, the net charge dictated the degree of mCherry colocalization with GFP(−6)-H_2_ in condensates. When both proteins were equivalent in net charge, they exhibited almost identical partitioning, localization, and condensate prevalence among a given cell population. Other properties such as the overall reversibility of the multi-protein condensate could be attributed to GFP(−6)-H_2_, which demonstrated condensate reversibility in single-protein condensates, although to a lesser extent. Overall, the same sequence-determinants governing single-protein systems could be extended to multi-protein systems when electrostatic interactions act as the main contributors to multivalent interactions. In the context of cell biology, these results highlight how spatiotemporal order at the cellular and subcellular scales can emerge from non-specific interactions at the molecular scale. We envision that a comprehensive understanding of sequence-determinants from both globular and disordered protein domains will help shed light on the functional significance of bacterial condensates in key cellular processes.

## Limitations of the Study

Results from our studies have focused on examining the sequence-encoded effects of net charge on the formation of bacterial condensates. These insights provide a foundation for further probing the dynamics, composition, and material properties of protein-based condensates. To study the dynamics in greater detail and overcome the resolution limitations of traditional fluorescence microscopy, super-resolution microscopy techniques such as photo-activated localization microscopy (PALM) can be used to probe temporal and spatial arrangement of condensates in live-cell imaging studies. In addition, the material properties of condensates in cells can be further characterized using single molecule localization microscopy (SMLM) to track the mobility of individual molecules in condensates and the cell cytoplasm. Determination of apparent diffusion coefficients will allow us to understand the material states of condensates.

From this work, we identified an inverse relationship between the charge density of disordered peptides and protein expression levels. Additional studies systematically varying the protein expression levels can be performed to explore the concentration-dependencies of intracellular phase separation. Protein concentrations in both the dilute and dense phases can be estimated by fluorescence microscopy to map phase diagrams in living cells and determine if the threshold concentration for phase separation, *c*_*sat*_, decreases as the net charge or valence of proteins increase.

## ACKNOWLEDGEMENTS

This work was supported by the National Science Foundation (DMR: 1848388) (J.L., V.Y., A.C.O.). J.L. also acknowledges support from the Blavatnik Fund for Engineering Innovations in Health.

## AUTHOR CONTRIBUTIONS

Conceptualization, J.L., V.Y., and A.C.O.; Investigation, J.L., and V.Y.; Data analysis, writing original draft, review and editing, J.L., V.Y., and A.C.O.

## STAR METHODS

### KEY RESOURCES TABLE

| REAGENT or RESOURCE | SOURCE | IDENTIFIER |
| --- | --- | --- |
| **Bacterial and virus strains** |  |  |
| NEB 5-alpha Competent *E. coli* (High Efficiency) | New England Biolabs | Cat#C2987H |
| NiCo21(DE3) Competent *E. coli* | New England Biolabs | Cat#C2529H |
| **Chemicals, peptides, and recombinant proteins** |  |  |
| Yeast Extract, Ultrapure | Alfa Aesar, Thermo Fisher Scientific | Cat#J23547-A1 |
| Enzymatic Digest Casein Hydrolysate | Thermo Fisher Scientific | Cat#AAJ12855P5 |
| Sodium Chloride (Crystalline/Certified ACS) | Thermo Fisher Scientific | Cat#S2713 |
| Bacto Agar | Becton Dickinson | Cat#214010 |
| Ampicillin (Sodium), USP Grade | GoldBio | A-301-5 |
| QIAprep Spin Miniprep Kit | Qiagen | Cat#27106 |
| Phusion High-Fidelity DNA Polymerase | New England Biolabs | Cat#M0530L |
| NcoI-HF | New England Biolabs | Cat#R3193S |
| XhoI | New England Biolabs | Cat#R0146S |
| XbaI | New England Biolabs | Cat#R0145S |
| DpnI | New England Biolabs | Cat#R0176S |
| TopVision Agarose | Thermo Fisher Scientific | Cat#R0492 |
| QIAquick Gel Extraction Kit | Qiagen | Cat#28706X4 |
| QIAquick PCR Purification Kit | Qiagen | Cat#28106 |
| T4 DNA Ligase | New England Biolabs | Cat#M0202S |
| NEBuilder HiFi DNA Assembly Master Mix | New England Biolabs | Cat#E2621L |
| Isopropyl-beta-D-thiogalactoside (IPTG) | GoldBio | Cat#I2481C |
| HisPur Ni-NTA Resin | Thermo Fisher Scientific | Cat#88221 |
| Sodium Phosphate, Monobasic (NaH<sub>2</sub>PO<sub>4</sub>) | Thermo Fisher Scientific | Cat#AC448170010 |
| Imidazole | Sigma-Aldrich | Cat#I202 |
| HEPES-NaOH | Thermo Fisher Scientific | Cat#BP210-500 |
| RNase A | Roche | Cat#10109142001 |
| DNase I, Bovine Pancreas | GoldBio | Cat#D-300-100 |
| SimplyBlue SafeStain | Thermo Fisher Scientific | Cat#LC6060 |
| MOPS-SDS Running Buffer (20X) | Thermo Fisher Scientific | Cat#J62847.AP |
| NuPAGE LDS Sample Buffer (4X) | Thermo Fisher Scientific | Cat#NP0007 |
| Ribonucleic Acid from Torula Yeast, Type VI | Sigma-Aldrich | Cat#R6625-25G |
| **Deposited data** |  |  |
| Raw microscopy data | This paper | Figshare<br>(https://doi.org/10.6084/m9.figshare.23289650) |
| **Oligonucleotides** |  |  |
| ssDNA oligos (see Table S1 for sequences) | Integrated DNA Technologies | N/A |
| **Recombinant DNA** |  |  |
| pET-6xHis-sfGFP | Yeong et al. (32) | Addgene Plasmid #199171 |
| pET-6xHis-GFP(0) | Yeong et al. (32) | Addgene Plasmid #199166 |
| pET-6xHis-GFP(+6) | Yeong et al. (32) | Addgene Plasmid #199160 |
| pET-6xHis-GFP(-6)-L<sub>1</sub> | This paper | Addgene Plasmid #205027 |
| pET-6xHis-GFP(-6)-L<sub>2</sub> | This paper | Addgene Plasmid #205028 |
| pET-6xHis-GFP(-6)-L<sub>3</sub> | This paper | Addgene Plasmid #205029 |
| pET-6xHis-GFP(-6)-H<sub>1</sub> | This paper | Addgene Plasmid #205030 |
| pET-6xHis-GFP(-6)-H<sub>2</sub> | This paper | Addgene Plasmid #205031 |
| pET-6xHis-GFP(-6)-H<sub>3</sub> | This paper | Addgene Plasmid #205032 |
| pET-6xHis-GFP(0)-L<sub>1</sub> | This paper | Addgene Plasmid #205033 |
| pET-6xHis-GFP(0)-L<sub>2</sub> | This paper | Addgene Plasmid #205034 |
| pET-6xHis-GFP(0)-L<sub>3</sub> | This paper | Addgene Plasmid #205035 |
| pET-6xHis-GFP(0)-H<sub>1</sub> | This paper | Addgene Plasmid #205036 |
| pET-6xHis-GFP(0)-H<sub>2</sub> | This paper | Addgene Plasmid #205037 |
| pET-6xHis-GFP(0)-H<sub>3</sub> | This paper | Addgene Plasmid #205038 |
| pET-6xHis-GFP(+6)-L<sub>1</sub> | This paper | Addgene Plasmid #205039 |
| pET-6xHis-GFP(+6)-L<sub>2</sub> | This paper | Addgene Plasmid #205040 |
| pET-6xHis-GFP(+6)-L<sub>3</sub> | This paper | Addgene Plasmid #205041 |
| pET-6xHis-GFP(+6)-H<sub>1</sub> | This paper | Addgene Plasmid #205042 |
| pET-6xHis-GFP(+6)-H<sub>2</sub> | This paper | Addgene Plasmid #205043 |
| pET-6xHis-GFP(+6)-H<sub>3</sub> | This paper | Addgene Plasmid #205044 |
| pET-6xHis-mCherry(-6) | This paper | Addgene Plasmid #205045 |
| pET-6xHis-mCherry(-6)-L<sub>1</sub> | This paper | Addgene Plasmid #205046 |
| pET-6xHis-mCherry(-6)-L<sub>2</sub> | This paper | Addgene Plasmid #205047 |
| pET-6xHis-mCherry(-6)-L<sub>3</sub> | This paper | Addgene Plasmid #205048 |
| pET-6xHis-GFP(-6)-H<sub>2</sub>_mCherry(-6) | This paper | Addgene Plasmid #205049 |
| pET-6xHis-GFP(-6)-H<sub>2</sub>_mCherry(-6)-L<sub>1</sub> | This paper | Addgene Plasmid #205050 |
| pET-6xHis-GFP(-6)-H<sub>2</sub>_mCherry(-6)-L<sub>2</sub> | This paper | Addgene Plasmid #205051 |
| pET-6xHis-GFP(-6)-H<sub>2</sub>_mCherry(-6)-L<sub>3</sub> | This paper | Addgene Plasmid #205052 |
| **Software and algorithms** |  |  |
| PyMOL (version 2.5.5) | Schrodinger LLC | https://pymol.org/2/ |
| FIJI ImageJ (version 2.9.0/1.53t) | NIH | https://imagej.net/software/fiji/ |
| MicrobeJ Plugin (version 5.13o(10)) | Ducret et al. (45) | https://www.microbej.com/download-2/ |
| MATLAB (R2022a) | Mathworks Inc. | https://www.mathworks.com/ |
| Prism (version 9.5.1) | GraphPad | https://www.graphpad.com/ |
| Adobe Illustrator (2022) | Adobe | https://www.adobe.com/products/illustrator.html |
| **Other** |  |  |
| High Performance Cover Glass 18x18 mm, no. 1.5 | Carl Zeiss | Cat#10474379 |
| Microscope Slides | Thermo Fisher Scientific | Cat#12-544-1 |
| Breathe-Easy Sealing Films | Diversified Biotech | Cat#BEM-1 |
| Costar Clear Multiple Well Plates | Corning | Cat#07-201-590 |
| Nalgene Sterile Syringe Filters, SFCA, 0.2 µm | Thermo Fisher Scientific | Cat#09-740-105 |
| Amicon™ Ultra-15 Centrifugal Filter | Sigma-Aldrich | Cat#UFC905008 |
| Bolt™ 4-12% Bis-Tris Plus gel | Thermo Fisher Scientific | Cat#NW04125BOX |
| Half Area Well Clear Flat Bottom TC-treated Microplate | Corning | Cat#3697 |

## RESOURCE AVAILABILITY

### Lead contact

Further information and requests for resources and reagents should be directed to and will be fulfilled by the lead contact, Allie Obermeyer.

### Materials availability

Plasmids generated in this study have been deposited to Addgene (#205027 – #205052).

### Data and code availability

Raw microscopy images are deposited on Figshare (doi: 10.6084/m9.figshare.23289650) and custom MATLAB codes are publicly available on GitHub (https://github.com/Obermeyer-Group) upon publication.

## EXPERIMENTAL MODEL AND SUBJECT DETAILS

### Plasmid Construction

The two different charge density cationic peptides were designed by repeating the base peptide motif, GRRGKKSRK or KRRRKK, one to three times for the L and H peptides, respectively. These peptides were appended to superfolder GFP (sfGFP) variants or mCherry using a combination of PCR, restriction enzyme digestion, and T4 ligation. An additional lysine residue was included between the C-terminus of sfGFP and the disordered peptide domain to establish charge equivalency between sfGFP and mCherry. All variants contained an N-terminal 6xHis tag with either a GGA or GG linker for sfGFP and mCherry, respectively. Primers (Integrated DNA Technologies) with engineered restriction enzyme sites were used to clone cationic peptides onto the C-terminus of sfGFP. For the sfGFP variants, the forward and reverse primers contained NcoI and XhoI cut sites, respectively. For the mCherry variants, the forward and reverse primers contained XbaI and XhoI cut sites, respectively. Primers were diluted to 10 μM in Milli-Q water. PCR reactions were performed using Phusion polymerase as instructed by New England Biolabs (NEB). Purified PCR products and the pETDuet vector backbone were digested with restriction enzymes, NcoI and XhoI for the sfGFP variants, or XbaI and XhoI for the mCherry variants. DpnI was added to the digestion reaction of PCR amplified sfGFP inserts. Reactions were run on a 1% agarose gel (TopVision agarose) and bands were excised and purified using a QIAquick gel extraction kit (Qiagen). The final plasmid was assembled using T4 ligase (NEB) using a 5:1 molar ratio of insert to vector for the GFP variants and a 3:1 molar ratio of insert to vector for the mCherry variants. 2 μL of the ligation reaction was used to transform chemically competent NEB5α cells. Following sequence verification by Genewiz, plasmids were transformed into NiCo21(DE3) cells (NEB). The bicistronic GFP(−6)-H_2_+mCherry-L_n_ variant was constructed using HiFi Assembly. Briefly, PCR reactions were performed to add overlapping regions. PCR products were purified using the QIAquick PCR purification kit (Qiagen). Purified fragments were assembled using NEBuilder HiFi DNA Assembly Master Mix and annealed at 50 °C for 15 min to insert the mCherry gene downstream of GFP. The assembled plasmid was transformed into NEB5α cells. After sequence verification by Oxford Nanopore sequencing (Plasmidsaurus), the plasmid was transformed into NiCo21(DE3) cells.

### Strains and Growth Conditions

Glycerol stocks of NiCo21(DE3) cells (NEB) transformed with each variant were streaked onto LB agar plates with added ampicillin (100 μg/mL, Gold Biotechnology). A single colony was inoculated into sterilized LB media supplemented with ampicillin (100 μg/mL) and grown in an incubator (Thermo Fisher Scientific MaxQ 6000) for 16-18 h at 37 °C with shaking at 225 rpm. The overnight cultures were back diluted to OD_600_ ∼ 0.1 in 25 mL LB media supplemented with ampicillin (100 μg/mL) in sterile 125 mL Erlenmeyer flasks. Cultures were grown at 37 °C, shaking at 225 rpm for 2-3 h. At OD_600_ = 0.7-1.0, cultures were induced with 1 mM isopropyl β-D-1-thiogalactopyranoside (IPTG, Gold Biotechnology) and then kept at 25 °C with shaking at 225 rpm for 24 h. 20 μL aliquots were taken from the cultures at various time points after induction for imaging via optical microscopy.

## METHOD DETAILS

### Cell Growth and Protein Expression Assays

The cellular growth of NiCo21(DE3) *E. coli* and fluorescent protein production was monitored for the GFP variants studied here using a plate-based growth assay. Three colonies containing the plasmid for each variant were selected from a freshly grown LB agar plate and grown to saturation overnight at 37 °C with shaking at 225 rpm in 1 mL LB supplemented with ampicillin in a 24-well plate. The optical density (OD_600_) was measured and each replicate was back-diluted to an OD_600_ of ∼ 0.1 with LB with added ampicillin, for a total volume of 1 mL per culture. Three wells of 1 mL LB were included to measure the background media fluorescence. After 2-3 h, when the cultures reached a corrected OD_600_ between 0.7-1.0, the cultures were induced with 1 mM IPTG. The plate was then incubated at 25 °C. At 0, 2, 4, 6, 8, 10, 20, and 24 h post-induction, the OD_600_ and GFP fluorescence (λ_ex_= 488 nm, λ_em_= 530 nm) were measured on an Infinite M200 Pro microplate reader (Tecan). The data were analyzed by dividing the background-corrected GFP fluorescence values by the background-corrected OD_600_ values. For variants co-expressing mCherry, the OD at 640 nm was measured instead to avoid interference with the protein absorbance. Timepoint measurements were taken at 0, 1, 2, 4, 6, 8, 19, and 24 h for the single-protein controls and at 2, 4, 6, 8, 19, and 24 h for the co-expressed variants. To account for the instrument bandwidth and ensure minimal crosstalk between proteins, GFP fluorescence was measured at λ_ex_= 470 and λ_em_= 510 and mCherry fluorescence was measured at λ_ex_= 570 and λ_em_= 610.

### Fluorescence and Optical Microscopy

Cell samples were applied to agarose pads for imaging. The agarose pads were made by preparing 1 w/v% agarose (TopVision) in Milli-Q water. 50 μL of melted agarose was then pipetted onto a 25 mm x 75 mm microscope slide and immediately covered by an 18 mm x 18 mm #1.5 coverslip (Thermo Fisher Scientific). After the agarose solidified, the coverslip was slowly removed and 2 μL of cell culture from the 20 μL aliquot was added on top of the agarose pad. The agarose pad with the sample was gently covered with the coverslip and sealed on all four sides with clear nail polish. Cells were imaged on agarose pads at 2, 8, and 24 h post-induction, or 2, 3, 4, 5, 6, 8, 9, 10, 12, and 24 h post-induction for the time course experiment (Figure 2). Images were taken using a 100X oil 1.40 NA UPlanSAPO objective (Olympus) with illumination by the EVOS GFP light cube (λ_ex_ = 470/22 nm; λ_em_ = 525/50 nm), EVOS Texas Red light cube (λ_ex_ = 585/29 nm; λ_em_ = 628/32 nm), and brightfield channels on an EVOS FL Auto 2 inverted fluorescent microscope. Approximately 10 fields of view were taken for each sample to acquire images with a sufficient number of cells per strain (>96 cells per biological replicate). At least 3 biological replicates were performed for each strain at each time point.

### Protein Purification for *In Vitro* State Diagrams

An overnight culture was inoculated into 1 L of LB media supplemented with ampicillin (100 μg/mL). The same growth and protein expression conditions as described above were used. Following protein expression, cells were harvested by centrifugation (Thermo Fisher Scientific Sorvall Legend XTR) in a swinging bucket rotor (Thermo Fisher Scientific TX-750) at 4,000 rpm for 15 min and resuspended in lysis buffer (50 mM NaH_2_PO_4_, 1 M NaCl, pH 8.0). The resuspended cell pellet was then subjected to one freeze thaw cycle. RNase A (225 μg/L of culture) and DNase I (200 μg/L of culture) were added to the thawed resuspended cells prior to lysis by sonication (cycle: 2 s on, 4 s off) at 40% amplitude using a 1/8 in probe for 10 min (Thermo Fisher Scientific). Soluble proteins were separated from cell debris by centrifugation in a fixed angle rotor (Thermo Fisher Scientific, Fiberlite F15-8×50cy) at 10,000 rpm for 30 min at room temperature. Proteins in the soluble fraction were collected and purified using immobilized metal affinity chromatography. Briefly, 10-15 mL of His-Pur Ni-NTA resin (Thermo Fisher Scientific) was used per L of cell culture. The resin was initially equilibrated in 3 column volumes of lysis buffer before incubation with the soluble cell lysate. Fractions of unbound lysate, wash (lysis buffer containing 50 mM imidazole) and elution (lysis buffer containing 250 mM imidazole) were collected. Fractions were analyzed on a Bolt 4-12% Bis-Tris Plus gel (Invitrogen) to determine protein purity. Samples were prepared in 4X LDS loading buffer and incubated at 95 °C for 10 min. 10 μL of each sample was run at 200 V for 22 min. Gels were stained following the SimplyBlue SafeStain protocol and imaged (S2C-E). Pure fractions were then dialyzed against a 50 mM HEPES-NaOH, pH 7.4 buffer for at least 3 h per equilibration with a total of 7 exchanges of buffer. Dialyzed protein samples were concentrated using Amicon Ultra centrifugal filter units with a 10 kDa molecular weight cutoff (Sigma-Aldrich).

### Turbidimetry Assay for *In Vitro* State Diagrams

Protein concentrations were obtained by measuring absorbance at 488 nm on a Cary 60 UV-Vis spectrophotometer (Agilent Technologies) and calculating concentrations using Beer’s Law. The extinction coefficient for superfolder GFP (ε = 8.33 × 10^5^ M^-1^ cm^-1^) was used to calculate the concentration for all supercharged GFP variants. Stock solutions of fluorescent protein (20 mg/mL and 0.2 mg/mL) and total RNA from torula yeast type VI (2 mg/mL and 0.02 mg/mL) (Sigma-Aldrich #R6625-25G) were prepared in 50 mM HEPES-NaOH, pH 7.4. The pH of the RNA stock was adjusted to pH 7.4 with 10 M NaOH immediately after dissolution. Protein and RNA stock solutions were filtered using a 0.22 μm SFCA Thermo Fisher Scientific Nalgene 25 mm syringe filter. State diagrams were constructed by mixing buffer, protein, and RNA at various mass ratios to a total volume of 50 μL in a tissue culture-treated polystyrene 96 well half area plate (Corning, #3697). The absorbance (A) of each sample was measured at 600 nm immediately after mixing on a plate reader (Tecan Infinite M200 Pro) at 25 °C and converted to turbidity using the following equation: Turbidity = 100 - 10^2-A^. Heatmaps of the turbidity values at each of the tested macromolecule concentrations were generated in MATLAB. For GFP(−6)-L_2_, a filled contour plot of turbidity isolines was also generated in MATLAB.

## QUANTIFICATION AND STATISTICAL ANALYSIS

### Electrostatic Maps of GFP Variants

The PDB file for sfGFP (PDB ID: 2B3P) was obtained from the RCSB protein data bank and loaded in PyMOL v2.5.5 (Schrödinger, LLC). Using the PDB file for sfGFP as a template, GFP(0) and GFP(+6) were made by introducing residue substitutions in PyMOL (32). PDBs of the minimized peptide structures were obtained using PEP-FOLD3 (46). The C-terminus of the globular domain was appended to the N-terminus of the disordered peptides in PyMOL and electrostatic surface maps were generated using the ABPS Electrostatics plugin.

### Image Pre-processing and Cell Segmentation

Fluorescence microscopy images were pre-processed using Fiji with a plug-in. Briefly, images were compiled into stacks and were subjected to a rolling ball background subtraction with a 100-pixel radius. Stacks of images were individually thresholded using the Li method in MicrobeJ and cells that met specific area, length, width, circularity, and angularity constraints were identified and considered for analysis. The following constraints were used for all image stacks: area = 100 – 2000 pixels^2^, length = 20 – max pixels, width = 0 – 20 pixels, circularity = 0.25 – max; curvature = 0 – max; sinuosity = 0 – max; solidity = 0 – max; intensity = 0 – max. Bacteria were detected using the fit shape, rod-shaped mode. Images of individual cells identified by MicrobeJ were loaded into MATLAB for processing and identification of condensates. Cell attributes including the cell area, mean width, and length were also imported from MicrobeJ. A custom MATLAB script was used to identify condensates and determine the fraction of condensate-containing cells, condensate area fraction, condensate intensity, and condensate localization.

### Condensate Thresholding within Cells

The 2D interpolated cell arrays from MicrobeJ were stored. Any remaining background pixels in the 2D array with an intensity lower than the minimum intensity from the cell contour were removed. The pixel intensity was sampled at three locations along the medial axis of each cell to approximate the intensity of a non-condensate area within the cell. In each of the three locations, the intensities of nine adjacent pixels were averaged and the lowest of the three averages was taken as the approximate average pixel intensity of the cytoplasm. All regions in a given cell with 20% higher intensity than the cell’s approximated average intensity and with at least 5 contiguous pixels at this higher intensity were classified as condensates. All other pixels were classified as the cytoplasm. Pixels from the cytoplasm and identified condensates were stored as 2D arrays along with the parent cell.

### Fraction of Cells with Condensates

All cells were classified as condensate-forming cells or non-condensate-forming cells. The fraction of condensate-containing cells for each replicate was calculated by dividing the number of condensate-forming cells by the total number of cells.

### Cell and Condensate Dimensions

Cell dimensions, including the width, length, and area were obtained from MicrobeJ. The area of each condensate within a cell was determined by MATLAB. Condensates were analyzed separately for cells containing multiple condensates. The area fraction of each condensate was calculated by dividing the area of the condensate by the area of the cell. For cells with multiple condensates, the area fractions occupied by all condensates were summed to obtain the total area fraction of condensates per cell.

### Cell and Condensate Pixel Intensities

The mean cell intensity within each cell contour was obtained from MicrobeJ. The mean intensities of the condensate and cytoplasm for each cell were calculated with MATLAB. For cells containing multiple condensates, the intensity was calculated separately for each condensate. The intensity ratio was calculated as the mean intensity of the condensate divided by the mean intensity of the cytoplasm. Data from MATLAB was transferred to GraphPad Prism (v. 9.5.1) for plotting and visualization.

### Condensate Localization Heat Maps

Localization heat maps were generated by binning the normalized cell position of the 1-pixel maxima from all condensates into a 20 by 10 array. Cells containing one condensate were aligned with the condensate in the positive end of the medial axis. A representative cell contour was generated by binning background pixels into a 20 by 10 array and generating a contour that represents 90% of all cells.

## References

1. Tetsuro Hirose, Kensuke Ninomiya, Shinichi Nakagawa, and Tomohiro Yamazaki. A guide to membraneless organelles and their various roles in gene regulation. Nature Reviews Molecular Cell Biology, 24(4):288–304, April 2023. ISSN 1471-0072, 1471-0080. doi: 10.1038/s41580-022-00558-8.

2. Andrew S. Lyon, William B. Peeples, and Michael K. Rosen. A framework for understanding the functions of biomolecular condensates across scales. Nature Reviews Molecular Cell Biology, 22(3):215–235, March 2021. ISSN 1471-0072, 1471-0080. doi: 10.1038/s41580-020-00303-z.

3. T Pederson. The plurifunctional nucleolus. Nucleic Acids Research, 26(17):3871–3876, September 1998. ISSN 13624962. doi: 10.1093/nar/26.17.3871.

4. Denis L. J. Lafontaine, Joshua A. Riback, Rümeyza Bascetin, and Clifford P. Brangwynne. The nucleolus as a multiphase liquid condensate. Nature Reviews Molecular Cell Biology, 22(3):165–182, March 2021. ISSN 1471-0072, 1471-0080. doi: 10.1038/s41580-020-0272-6.

5. S Ramón Cajal. Un sencillo metodo de coloracion seletiva del reticulo protoplasmatico y sus efectos en los diversos organos nerviosos de vertebrados e invertebrados. Trab. Lab. Invest. Biol.(Madrid), 2:129–221, 1903.

6. Noriko Saitoh, Chris S. Spahr, Scott D. Patterson, Paula Bubulya, Andrew F. Neuwald, and David L. Spector. Proteomic Analysis of Interchromatin Granule Clusters. Molecular Biology of the Cell, 15(8):3876–3890, August 2004. ISSN 1059-1524, 1939-4586. doi: 10.1091/mbc.e04-03-0253.

7. Archa H. Fox, Yun Wah Lam, Anthony K.L. Leung, Carol E. Lyon, Jens Andersen, Matthias Mann, and Angus I. Lamond. Paraspeckles. Current Biology, 12(1):13–25, January 2002. ISSN 09609822. doi: 10.1016/S0960-9822(01)00632-7.

8. Natalie Gilks, Nancy Kedersha, Maranatha Ayodele, Lily Shen, Georg Stoecklin, Laura M Dember, and Paul Anderson. Stress Granule Assembly Is Mediated by Prion-like Aggregation of TIA-1. Molecular Biology of the Cell, 15, 2004. doi: 10.1091/mbc.e04-08-0715.

9. Clifford P. Brangwynne, Christian R. Eckmann, David S. Courson, Agata Rybarska, Carsten Hoege, Jöbin Gharakhani, Frank Jülicher, and Anthony A. Hyman. Germline P Granules Are Liquid Droplets That Localize by Controlled Dissolution/Condensation. Science, 324(5935):1729–1732, June 2009. ISSN 0036-8075, 1095-9203. doi: 10.1126/science.1172046.

10. Zixu Gao, Wenchang Zhang, Runlei Chang, Susu Zhang, Guiwen Yang, and Guoyan Zhao. Liquid-Liquid Phase Separation: Unraveling the Enigma of Biomolecular Condensates in Microbial Cells. Frontiers in Microbiology, 12:751880, October 2021. ISSN 1664-302X. doi: 10.3389/fmicb.2021.751880.

11. Christopher A. Azaldegui, Anthony G. Vecchiarelli, and Julie S. Biteen. The emergence of phase separation as an organizing principle in bacteria. Biophysical Journal, 120(7):1123–1138, April 2021. ISSN 00063495. doi: 10.1016/j.bpj.2020.09.023.

12. Elio A. Abbondanzieri and Anne S. Meyer. More than just a phase: the search for membraneless organelles in the bacterial cytoplasm. Current Genetics, 65(3):691–694, June 2019. ISSN 0172-8083, 1432-0983. doi: 10.1007/s00294-018-00927-x.

13. Nisansala S. Muthunayake, Dylan T. Tomares, W. Seth Childers, and Jared M. Schrader. Phase-separated bacterial ribonucleoprotein bodies organize mRNA decay. WIREs RNA, 11(6), November 2020. ISSN 1757-7004, 1757-7012. doi: 10.1002/wrna.1599.

14. Chris Greening and Trevor Lithgow. Formation and function of bacterial organelles. Nature Reviews Microbiology, 18(12):677–689, December 2020. ISSN 1740-1526, 1740-1534. doi: 10.1038/s41579-020-0413-0.

15. Megan C. Cohan and Rohit V. Pappu. Making the Case for Disordered Proteins and Biomolecular Condensates in Bacteria. Trends in Biochemical Sciences, 45(8):668–680, August 2020. ISSN 09680004. doi: 10.1016/j.tibs.2020.04.011.

16. Salman F. Banani, Hyun O. Lee, Anthony A. Hyman, and Michael K. Rosen. Biomolecular condensates: organizers of cellular biochemistry. Nature Reviews Molecular Cell Biology, 18(5):285–298, May 2017. ISSN 1471-0072, 1471-0080. doi: 10.1038/nrm.2017.7.

17. M. E. Cates and T. A. Witten. Chain conformation and solubility of associating polymers. Macromolecules, 19(3):732–739, March 1986. ISSN 0024-9297, 1520-5835. doi: 10.1021/ma00157a042.

18. Alexander N. Semenov and Michael Rubinstein. Thermoreversible Gelation in Solutions of Associative Polymers. 1. Statics. Macromolecules, 31(4):1373–1385, February 1998. ISSN 0024-9297, 1520-5835. doi: 10.1021/ma970616h.

19. Garrett M. Ginell and Alex S. Holehouse. An Introduction to the Stickers-and-Spacers Framework as Applied to Biomolecular Condensates. In Huan-Xiang Zhou, Jan-Hendrik Spille, and Priya R. Banerjee, editors, Phase-Separated Biomolecular Condensates, volume 2563, pages 95–116. Springer US, New York, NY, 2023. ISBN 978-1-07-162662-7 978-1-07-162663-4. doi: 10.1007/978-1-0716-2663-4_4. Series Title: Methods in Molecular Biology.

20. Tanja Mittag and Rohit V. Pappu. A conceptual framework for understanding phase separation and addressing open questions and challenges. Molecular Cell, 82(12):2201–2214, June 2022. ISSN 10972765. doi: 10.1016/j.molcel.2022.05.018.

21. Benjamin S. Schuster, Ellen H. Reed, Ranganath Parthasarathy, Craig N. Jahnke, Reese M. Caldwell, Jessica G. Bermudez, Holly Ramage, Matthew C. Good, and Daniel A. Hammer. Controllable protein phase separation and modular recruitment to form responsive membraneless organelles. Nature Communications, 9(1):2985, July 2018. ISSN 2041-1723. doi: 10.1038/s41467-018-05403-1.

22. Joseph R. Simon, Nick J. Carroll, Michael Rubinstein, Ashutosh Chilkoti, and Gabriel P. López. Programming molecular self-assembly of intrinsically disordered proteins containing sequences of low complexity. Nature Chemistry, 9(6):509–515, June 2017. ISSN 1755-4330, 1755-4349. doi: 10.1038/nchem.2715.

23. Robin Van Der Lee, Marija Buljan, Benjamin Lang, Robert J. Weatheritt, Gary W. Daughdrill, A. Keith Dunker, Monika Fuxreiter, Julian Gough, Joerg Gsponer, David T. Jones, Philip M. Kim, Richard W. Kriwacki, Christopher J. Oldfield, Rohit V. Pappu, Peter Tompa, Vladimir N. Uversky, Peter E. Wright, and M. Madan Babu. Classification of Intrinsically Disordered Regions and Proteins. Chemical Reviews, 114(13):6589–6631, July 2014. ISSN 0009-2665, 1520-6890. doi: 10.1021/cr400525m.

24. Begoña Monterroso, Silvia Zorrilla, Marta Sobrinos-Sanguino, Miguel A Robles-Ramos, Marina López-Álvarez, William Margolin, Christine D Keating, and Germán Rivas. Bacterial FtsZ protein forms phase-separated condensates with its nucleoid-associated inhibitor SlmA. EMBO reports, 20(1):e45946, January 2019. ISSN 1469-221X, 1469-3178. doi: 10.15252/embr.201845946.

25. Min Kyung Shinn, Megan C. Cohan, Jessie L. Bullock, Kiersten M. Ruff, Petra A. Levin, and Rohit V. Pappu. Connecting sequence features within the disordered C-terminal linker of Bacillus subtilis FtsZ to functions and bacterial cell division. Proceedings of the National Academy of Sciences, 119(42):e2211178119, October 2022. ISSN 0027-8424, 1091-6490. doi: 10.1073/pnas.2211178119.

26. Nadra Al-Husini, Dylan T. Tomares, Obaidah Bitar, W. Seth Childers, and Jared M. Schrader. α-Proteobacterial RNA Degradosomes Assemble Liquid-Liquid Phase-Separated RNP Bodies. Molecular Cell, 71(6):1027–1039.e14, September 2018. ISSN 10972765. doi: 10.1016/j.molcel.2018.08.003.

27. Nadra Al-Husini, Dylan T. Tomares, Zechariah J. Pfaffenberger, Nisansala S. Muthunayake, Mohammad A. Samad, Tiancheng Zuo, Obaidah Bitar, James R. Aretakis, Mohammed-Husain M. Bharmal, Alisa Gega, Julie S. Biteen, W. Seth Childers, and Jared M. Schrader. BR-Bodies Provide Selectively Permeable Condensates that Stimulate mRNA Decay and Prevent Release of Decay Intermediates. Molecular Cell, 78(4):670–682.e8, May 2020. ISSN 10972765. doi: 10.1016/j.molcel.2020.04.001.

28. Xin Ge, Andrew J. Conley, Jim E. Brandle, Ray Truant, and Carlos D. M. Filipe. In Vivo Formation of Protein Based Aqueous Microcompartments. Journal of the American Chemical Society, 131(25):9094–9099, July 2009. ISSN 0002-7863, 1520-5126. doi: 10.1021/ja902890r.

29. Shao-Peng Wei, Zhi-Gang Qian, Chun-Fei Hu, Fang Pan, Meng-Ting Chen, Sang Yup Lee, and Xiao-Xia Xia. Formation and functionalization of membraneless compartments in Escherichia coli. Nature Chemical Biology, 16(10):1143–1148, October 2020. ISSN 1552-4450, 1552-4469. doi: 10.1038/s41589-020-0579-9.

30. Michael Dzuricky, Bradley A. Rogers, Abdulla Shahid, Paul S. Cremer, and Ashutosh Chilkoti. De novo engineering of intracellular condensates using artificial disordered proteins. Nature Chemistry, 12(9):814–825, September 2020. ISSN 1755-4330, 1755-4349. doi: 10.1038/s41557-020-0511-7.

31. Yifan Dai, Mina Farag, Dongheon Lee, Xiangze Zeng, Kyeri Kim, Hye-in Son, Xiao Guo, Jonathan Su, Nikhil Peterson, Javid Mohammed, Max Ney, Daniel Mark Shapiro, Rohit V. Pappu, Ashutosh Chilkoti, and Lingchong You. Programmable synthetic biomolecular condensates for cellular control. Nature Chemical Biology, 19(4):518–528, April 2023. ISSN 1552-4450, 1552-4469. doi: 10.1038/s41589-022-01252-8.

32. Vivian Yeong, Emily G. Werth, Lewis M. Brown, and Allie C. Obermeyer. Formation of Biomolecular Condensates in Bacteria by Tuning Protein Electrostatics. ACS Central Science, 6(12):2301–2310, December 2020. ISSN 2374-7943, 2374-7951. doi: 10.1021/acscentsci.0c01146.

33. Vivian Yeong, Jou-wen Wang, Justin M. Horn, and Allie C. Obermeyer. Intracellular phase separation of globular proteins facilitated by short cationic peptides. Nature Communications, 13(1):7882, December 2022. ISSN 2041-1723. doi: 10.1038/s41467-022-35529-2.

34. Jorge R. Espinosa, Jerelle A. Joseph, Ignacio Sanchez-Burgos, Adiran Garaizar, Daan Frenkel, and Rosana Collepardo-Guevara. Liquid network connectivity regulates the stability and composition of biomolecular condensates with many components. Proceedings of the National Academy of Sciences, 117(24):13238–13247, June 2020. ISSN 0027-8424, 1091-6490. doi: 10.1073/pnas.1917569117.

35. Yifan Dai, Lingchong You, and Ashutosh Chilkoti. Engineering synthetic biomolecular condensates. Nature Reviews Bioengineering, April 2023. ISSN 2731-6092. doi: 10.1038/s44222-023-00052-6.

36. Sara Molinari, David L. Shis, Shyam P. Bhakta, James Chappell, Oleg A. Igoshin, and Matthew R. Bennett. A synthetic system for asymmetric cell division in Escherichia coli. Nature Chemical Biology, 15(9):917–924, September 2019. ISSN 1552-4450, 1552-4469. doi: 10.1038/s41589-019-0339-x.

37. Keren Lasker, Lexy Von Diezmann, Xiaofeng Zhou, Daniel G. Ahrens, Thomas H. Mann, W. E. Moerner, and Lucy Shapiro. Selective sequestration of signalling proteins in a membraneless organelle reinforces the spatial regulation of asymmetry in Caulobacter crescentus. Nature Microbiology, 5(3):418–429, January 2020. ISSN 2058-5276. doi: 10.1038/s41564-019-0647-7.

38. Da-Wei Lin, Yang Liu, Yue-Qi Lee, Po-Jiun Yang, Chia-Tse Ho, Jui-Chung Hong, Jye-Chian Hsiao, Der-Chien Liao, An-Jou Liang, Tzu-Chiao Hung, Yu-Chuan Chen, Hsiung-Lin Tu, Chao-Ping Hsu, and Hsiao-Chun Huang. Construction of intracellular asymmetry and asymmetric division in Escherichia coli. Nature Communications, 12(1):888, February 2021. ISSN 2041-1723. doi: 10.1038/s41467-021-21135-1.

39. Mehmet U. Caglar, John R. Houser, Craig S. Barnhart, Daniel R. Boutz, Sean M. Carroll, Aurko Dasgupta, Walter F. Lenoir, Bartram L. Smith, Viswanadham Sridhara, Dariya K. Sydykova, Drew Vander Wood, Christopher J. Marx, Edward M. Marcotte, Jeffrey E. Barrick, and Claus O. Wilke. The E. coli molecular phenotype under different growth conditions. Scientific Reports, 7(1):45303, April 2017. ISSN 2045-2322. doi: 10.1038/srep45303.

40. Thomas Prossliner, Kristoffer Skovbo Winther, Michael Askvad Sørensen, and Kenn Gerdes. Ribosome Hibernation. Annual Review of Genetics, 52(1):321–348, November 2018. ISSN 0066-4197, 1545-2948. doi: 10.1146/annurev-genet-120215-035130.

41. D Mengin-Lecreulx and J van Heijenoort. Effect of growth conditions on peptidoglycan content and cytoplasmic steps of its biosynthesis in Escherichia coli. Journal of Bacteriology, 163(1):208–212, July 1985. ISSN 0021-9193, 1098-5530. doi: 10.1128/jb.163.1.208-212.1985.

42. Juana María Navarro Llorens, Antonio Tormo, and Esteban Martínez-García. Stationary phase in gram-negative bacteria. FEMS Microbiology Reviews, 34(4):476–495, July 2010. ISSN 1574-6976. doi: 10.1111/j.1574-6976.2010.00213.x.

43. William M. Aumiller, Fatma Pir Cakmak, Bradley W. Davis, and Christine D. Keating. RNA-Based Coacervates as a Model for Membraneless Organelles: Formation, Properties, and Interfacial Liposome Assembly. Langmuir, 32(39):10042–10053, October 2016. ISSN 0743-7463, 1520-5827. doi: 10.1021/acs.langmuir.6b02499.

44. Damian Wollny, Benjamin Vernot, Jie Wang, Maria Hondele, Aram Safrastyan, Franziska Aron, Julia Micheel, Zhisong He, Anthony Hyman, Karsten Weis, J. Gray Camp, T.-Y. Dora Tang, and Barbara Treutlein. Characterization of RNA content in individual phase-separated coacervate microdroplets. Nature Communications, 13(1):2626, May 2022. ISSN 2041-1723. doi: 10.1038/s41467-022-30158-1.

45. Adrien Ducret, Ellen M. Quardokus, and Yves V. Brun. MicrobeJ, a tool for high throughput bacterial cell detection and quantitative analysis. Nature Microbiology, 1(7):16077, June 2016. ISSN 2058-5276. doi: 10.1038/nmicrobiol.2016.77.

46. Alexis Lamiable, Pierre Thévenet, Julien Rey, Marek Vavrusa, Philippe Derreumaux, and Pierre Tufféry. PEP-FOLD3: faster de novo structure prediction for linear peptides in solution and in complex. Nucleic Acids Research, 44(W1):W449–W454, July 2016. ISSN 0305-1048, 1362-4962. doi: 10.1093/nar/gkw329.

