## Supporting Information for "Charge-Patterned Disordered Peptides Tune Intracellular Phase Separation in Bacteria"

**This PDF file includes:**

Figures S1 to S5
Table S1

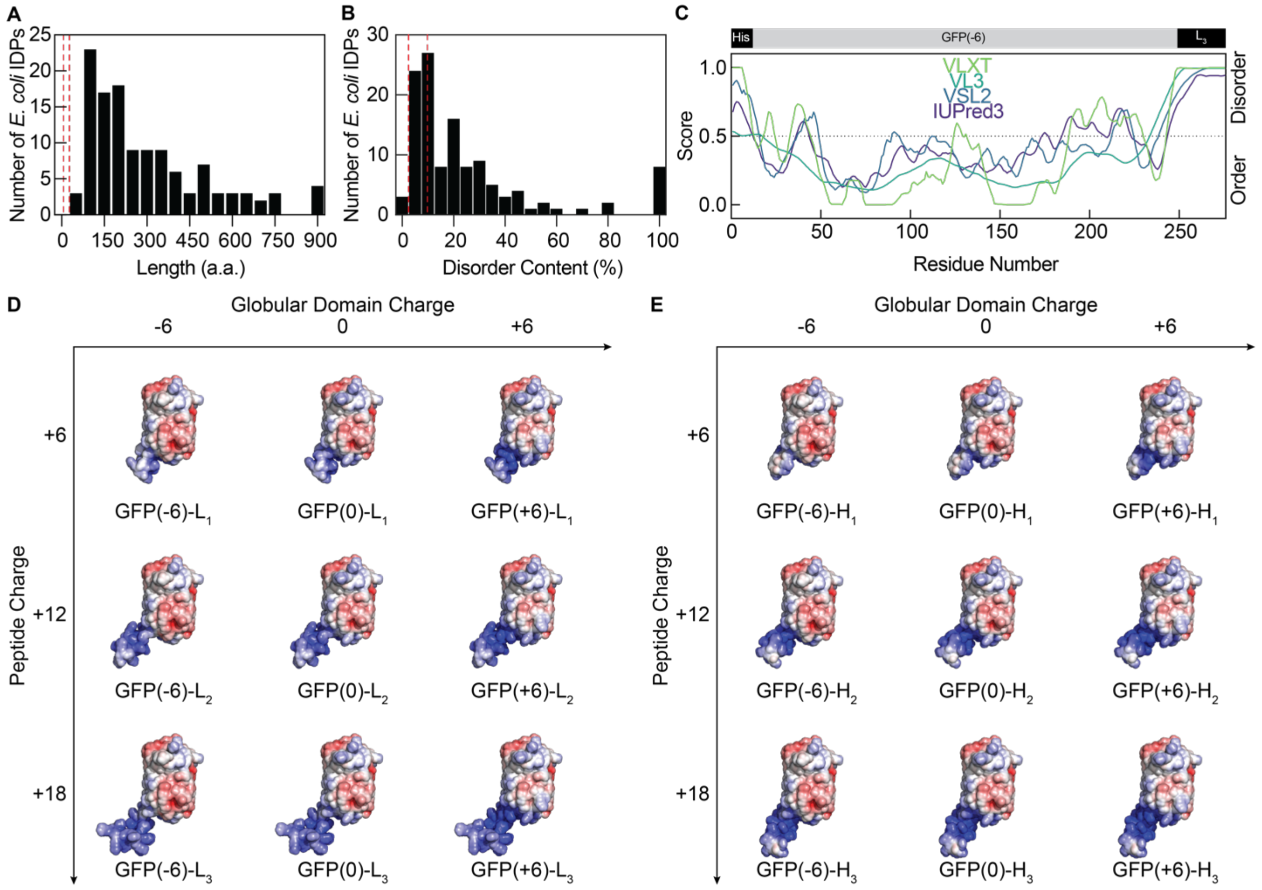


| Peptide | Charge | Sequence |
|---|---|---|
| $L_1$ | +6 | GRRGKKSRK |
| $L_2$ | +12 | GRRGKKSRK GRRGKKSRK |
| $L_3$ | +18 | GRRGKKSRK GRRGKKSRK GRRGKKSRK |

| Peptide | Charge | Sequence |
|---|---|---|
| $H_1$ | +6 | KRRRKK |
| $H_2$ | +12 | KRRRKK KRRRKK |
| $H_3$ | +18 | KRRRKK KRRRKK KRRRKK |

**Figure S1. Disorder and charge-patterning of C-terminal cationic peptides**

(A) Sequence lengths of endogenous IDPs identified from *E. coli* (K12) by DisProt[1–3]. Red dashed lines indicate the minimum and maximum lengths of engineered disordered cationic peptides.

(B) Disorder content of endogenous IDPs identified from *E. coli* (K12) by DisProt. Red dashed lines indicate the minimum and maximum disorder content of engineered disordered cationic peptides, approximated by dividing the length of the disordered peptide domain by the full length of the protein.

(C) Prediction of disorder scores by PONDR[4] and IUPred3[5] for a representative engineered protein variant, GFP(-6)-$L_3$.

(D) Electrostatic surface representations of the GFP variants with low charge density (L) peptides.

(E) Electrostatic surface representations of the GFP variants with high charge density (H) peptides.

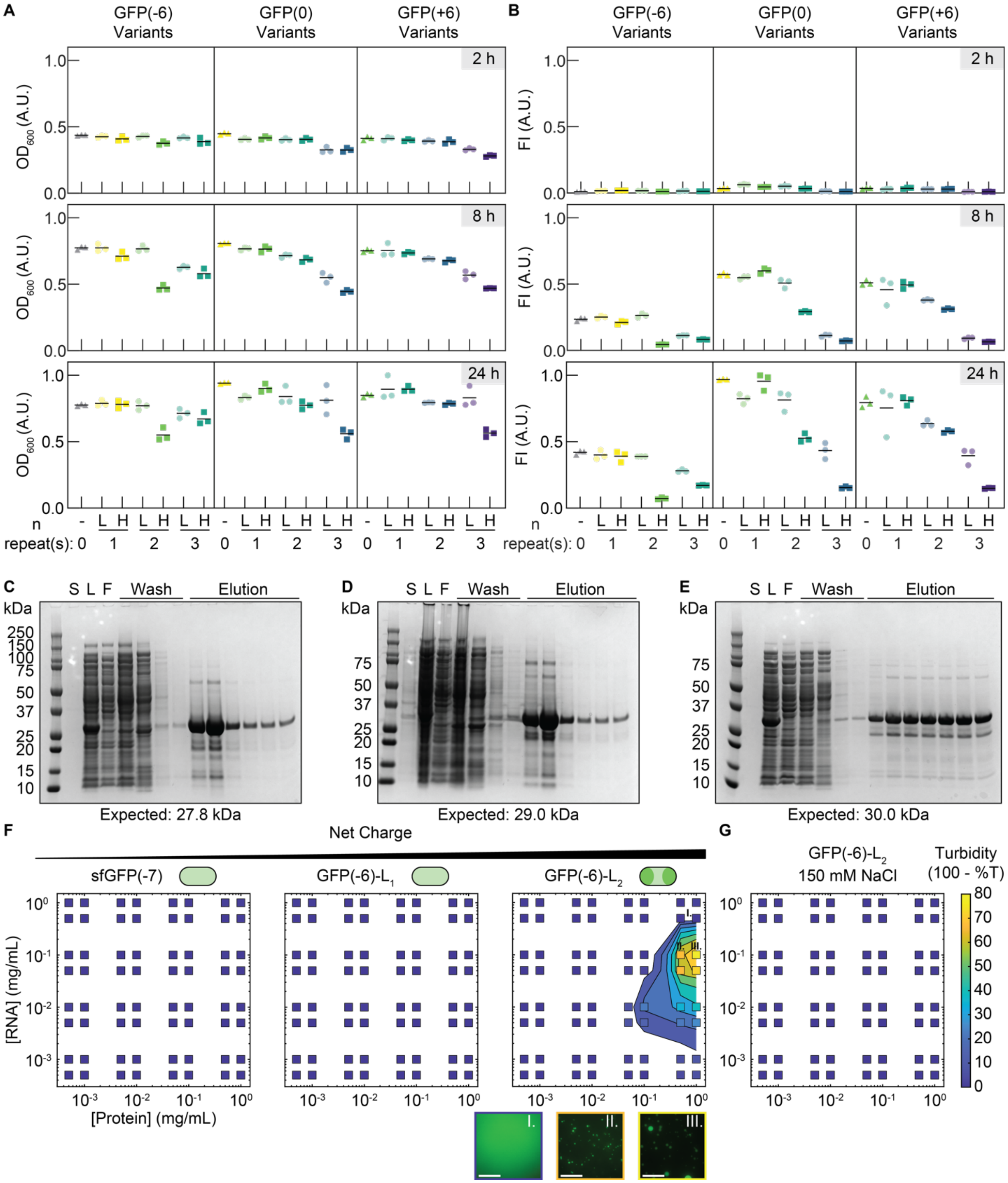


**Figure S2. Cell growth and protein expression decrease with increasing cationic peptide length.**

(A) Normalized $OD_{600}$ at 2, 8, and 24 h post-induction shows that both higher peptide charge density and increasing peptide length decrease cell growth. Data was normalized to GFP(+6)-$L_1$ at 24 h as the maximum.

(B) Normalized FI at 2, 8, and 24 h post-induction shows that both higher peptide charge density and increasing peptide length decrease protein expression. Data was normalized to GFP(0)-$H_1$ at 24 h as the maximum. For (A) and (B), three biological replicates and the respective means are shown. Triangles, circles, and squares indicate isotropic variants, low charge density peptide variants, and high charge density peptide variants, respectively.

(C) SDS-PAGE gel for the purification of sfGFP(-7).

(D) SDS-PAGE gel for the purification of GFP(-6)-$L_1$.

(E) SDS-PAGE gel for the purification of GFP(-6)-$L_2$. For (C-E), lanes depict the supernatant (S), lysate (L), flowthrough (F), washes, and elution fractions, in order.
(F) Phase diagrams as determined by turbidity measurements of purified GFP variants with total yeast RNA mixed at the indicated concentrations (boxes) in 50 mM HEPES-NaOH, pH 7.4. *In vitro* phase separation is dependent on the net charge of the protein scaffold. Shading within boxes depicts turbidity values, and filled regions represent turbidity contours. Fluorescence microscopy images of indicated mixtures. Scale bars are 25 μm.
(G) GFP(-6)-$L_2$ and RNA do not phase separate at the indicated concentrations in 50 mM HEPES-NaOH, 150 mM NaCl, pH 7.4, suggesting that the interactions are mainly electrostatic.

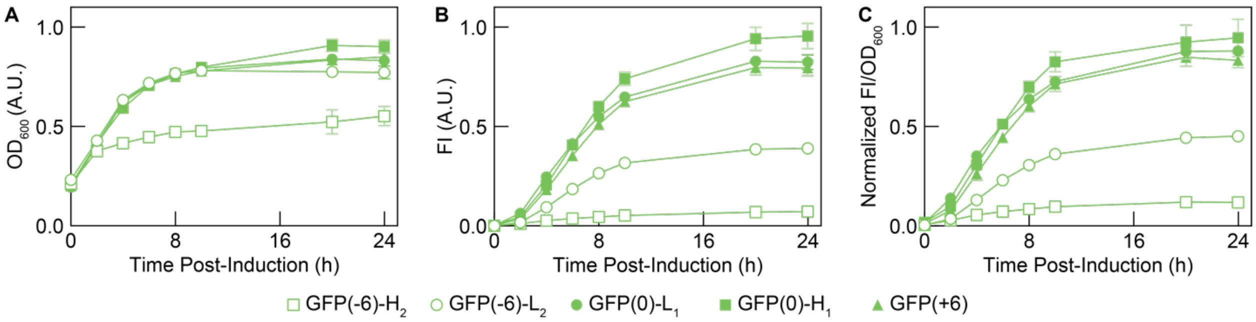


**Figure S3. Cell growth and protein expression over time for all +6 net charge variants.**

(A) Normalized $OD_{600}$ over time for all +6 net charge variants. GFP(-6)-$H_2$ has relatively lower cell growth than the other variants. Data was normalized to GFP(+6)-$L_1$ at 24 h (Figure S4) as the maximum.

(B) Normalized FI over time for all +6 net charge variants. Both GFP(-6) variants have relatively lower protein expression than the other variants. Data was normalized to GFP(0)-$H_1$ at 24 h as the maximum.

(C) Normalized FI/$OD_{600}$ over time for all +6 net charge variants. Data was normalized to GFP(0)-$H_1$ at 24 h as the maximum. For (A), (B), and (C), data points show the mean and error bars indicate the standard deviation of three biological replicates.

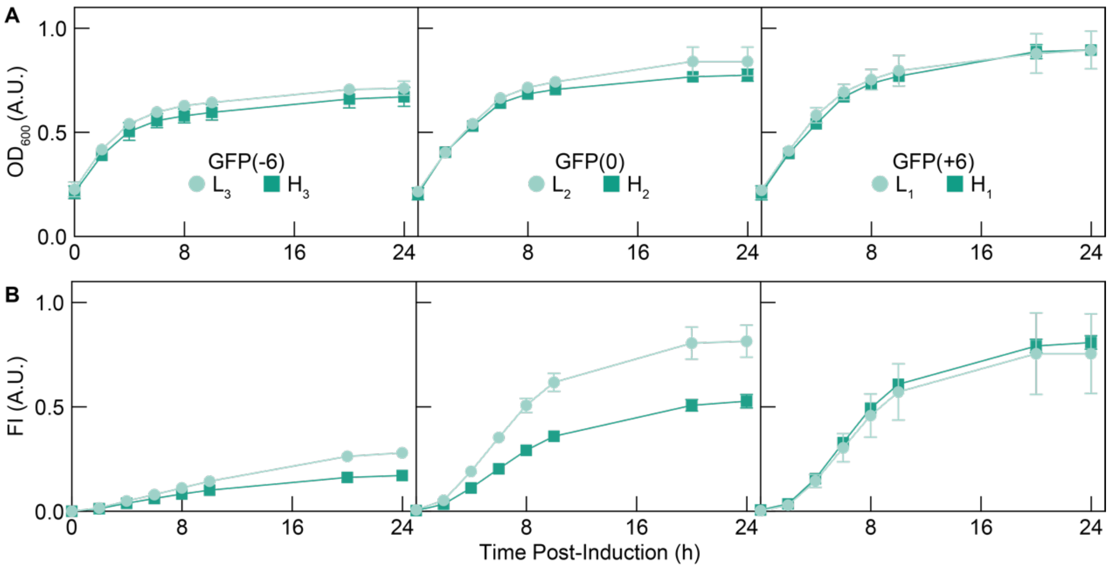


**Figure S4. Cell growth and protein expression over time for all +12 net charge variants.**
(A) Normalized $OD_{600}$ over time for all +12 net charge variants. Data was normalized to GFP(+6)-$L_1$ at 24 h as the maximum.
(B) Normalized FI over time for all +12 net charge variants. Both GFP(-6)-$H_3$ and GFP(0)-$H_2$ have relatively lower protein expression than GFP(-6)-$L_3$ and GFP(0)-$L_2$. Data was normalized to GFP(0)-$H_1$ at 24 h as the maximum. For (A) and (B), data points show the mean and error bars indicate the standard deviation of three biological replicates.

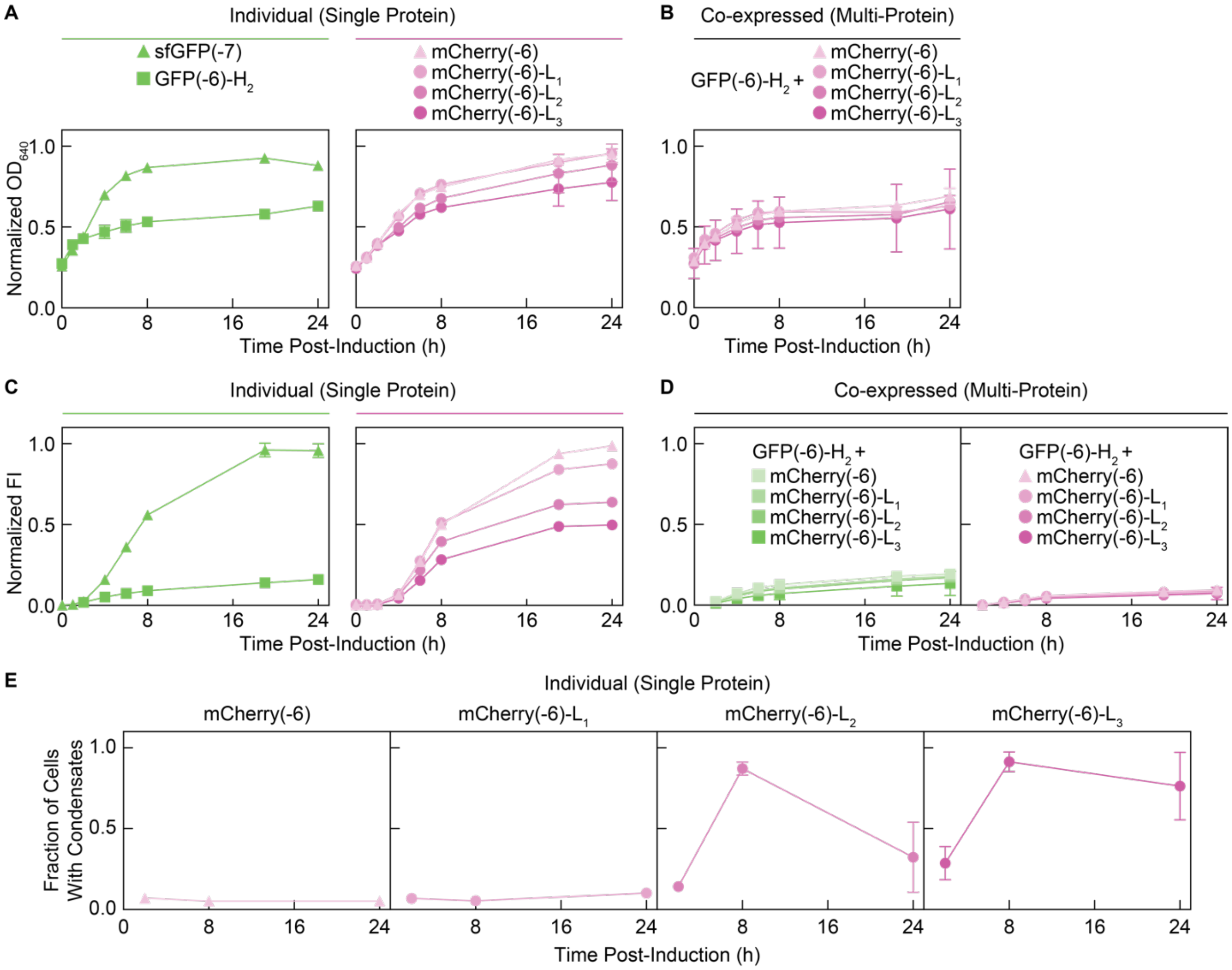


**Figure S5. Cell growth and protein expression over time for all GFP and mCherry co-expressed variants.**
(A) Normalized $OD_{600}$ over time for cells expressing a single protein shows that the peptide domain reduces cell growth. Data was normalized to mCherry(-6) at 24 h as the maximum.
(B) Normalized $OD_{600}$ over time for cells co-expressing GFP(-6)-$H_2$ and mCherry-$L_n$. Data was normalized to mCherry(-6) at 24 h as the maximum.
(C) Normalized FI over time for cells expressing a single protein shows that the peptide domain reduces protein expression. Data was normalized sfGFP(-7) at 19 h as the maximum.
(D) Normalized FI over time for cells co-expressing GFP(-6)-$H_2$ and mCherry-$L_n$. Data was normalized sfGFP(-7) at 19 h as the maximum. For (A), (B), (C), and (D), data points show the mean and error bars indicate the standard deviation of three biological replicates.
(E) A net charge of +6 is required for the phase separation of mCherry-$L_n$ variants when expressed alone.

**Table S1. Primer sequences for the construction of L- and H-peptide variants**

| Plasmid | DNA Template | Primer | Primer Sequence (5' to 3') |
|---|---|---|---|
| **GFP Variants** | | | |
| GFP(-6)-L1 | sfGFP | Fwd[a] | GATATACCATGGGTCATCACCACCACC |
| | | Rev[b] | TAATCTCGAG/TTA/CTTTCGAGATTTCTTGCCTCTTC GTCCTTT/CTTGTACAGCTCGTCCA |
| GFP(-6)-L2 | sfGFP | Rev | TAATCTCGAG/TTATTA/CTTTCGAGATTTCTTGCCTC TTCGTCCCTTTCGAGATTTCTTGCCTCTTCGTCCTTT/ CTTGTACAGCTCGTCCA |
| GFP(-6)-L3 | GFP(-6)-L1 | Rev | ATACTCGAG/TTATTA/CTTCCGCGATTTCTTCCCTCT TCGACCCTTCCGCGATTTCTTCCCTCTTCGACC/CTT TCGAGATTTCTTGCCTC |
| GFP(-6)-H1 | sfGFP | Rev | TAATCTCGAG/TTATTA/TTTCTTCCTTCTCCTCTTTT T/CTTGTACAGCTCGTCCATTC |
| GFP(-6)-H2 | sfGFP | Rev | TAATCTCGAG/TTA/TTTCTTCCTTCTCCTCTTTTTCTT CCTTCTCCTCTTTTT/CTTGTACAGCTCGTCCATTC |
| GFP(-6)-H3 | GFP(-6)-H1 | Rev | ATACTCGAG/TTATTA/TTTCTTGCGTCTACGCTTTTT CTTGCGTCTACGCTT/TTTCTTCCTTCTCCTCTTTT |
| GFP(0)-L1 | GFP(0) | Rev | ATACTCGAG/TTATTA/CTTCCGCGATTTCTTCCCTCT TCGACC/CTTGTAGCGTTCGTCCATTC |
| GFP(0)-L2 | GFP(0) | Rev | ATACTCGAG/TTATTA/CTTCCGCGATTTCTTCCCTCT TCGACCCTTCCGCGATTTCTTCCCTCTTCGACC/CTT GTAGCGTTCGTCCATTC |
| GFP(0)-L3 | GFP(0)-L2 | Rev | ATACTCGAG/TTATTA/CTTCCTAGATTTCTTACCCCT TCTTCC/CTTCCGCGATTTCTTCCCTCTTCGACCCTTC CGC |
| GFP(0)-H1 | GFP(0) | Rev | ATACTCGAG/TTATTA/TTTCTTGCGTCTACGCTT/CT TGTAGCGTTCGTCCATTC |
| GFP(0)-H2 | GFP(0) | Rev | ATACTCGAG/TTATTA/TTTCTTGCGTCTACGCTTTTT CTTGCGTCTACGCTT/CTTGTAGCGTTCGTCCATTC |
| GFP(0)-H3 | GFP(0)-H2 | Rev | ATACTCGAG/TTATTA/CTTTTTACGTCTTCGTTT/TTT CTTGCGTCTACGCTTTTTCTTG |
| GFP(+6)-L1 | GFP(+6) | Rev | Same as rev for GFP(0)-L1 |
| GFP(+6)-L2 | GFP(+6) | Rev | Same as rev for GFP(0)-L2 |
| GFP(+6)-L3 | GFP(+6)-L2 | Rev | ATACTCGAG/TTATTA/CTTCCTAGATTTCTTACCCCT TCTTCC/CTTCCGCGATTTCTTCCCTCTTCGACCCTTC CGC |
| GFP(+6)-H1 | GFP(+6) | Rev | Same as rev for GFP(0)-H1 |
| GFP(+6)-H2 | GFP(+6) | Rev | Same as rev for GFP(0)-H2 |
| GFP(+6)-H3 | GFP(+6)-H2 | Rev | Same as rev for GFP(0)-H3 |
| **mCherry Variants** | | | |
| mCherry(-6)-L1 | mCherry(-6) | Fwd[c] | CGGATAACAATTCCCCTCTAGA |
| | | Rev | TAATCTCGAG/TCATTA/CTTTCGAGATTTCTTGCCTC TTCGTCC/CTTGTACAGCTCGTCCA |
| mCherry(-6)-L2 | mCherry(-6)-L1 | Rev | TAATCTCGAG/TCATTA/CTTTCGAGATTTCTTGCCTC TTCGTCC/CTTTCGAGATTTCTTGC |
| mCherry(-6)-L3 | mCherry(-6)-L1 | Rev | ATACTCGAG/TTATTA/CTTCCGCGATTTCTTCCCTCT TCGACCCTTCCGCGATTTCTTCCCTCTTCGACC/CTT TCGAGATTTCTTGCCTC |
| **Bicistronic GFP+mCherry Variants** | | | |
| GFP(-6)-H2 + mCherry(-6) | GFP(-6)-H2, mCherry(-6) | mCherry_Fwd | TTAACTTTAAGAAGGAGATATACATATG |
| | | mCherry_Rev | TCATTACTTGTACAGCTCG |
| | | GFP_Fwd1 | ATATGGTGCACTCTCAGTACAATCTGCTC |

| | | | |
|---|---|---|---|
| | | GFP_Rev1 | TATCTCCTTCTTAAAGTTAATTATTTCTTCCTTCTCC TCTTTT |
| | | GFP_Fwd2 | ACGAGCTGTACAAGTAATGACTCGAGTCTGGTAAA GAAAC |
| | | GFP_Rev2 | GTACTGAGAGTGCACCATATATGCGGTG |
| GFP(-6)-H2 + mCherry(-6)-L1 | GFP(-6)-H2, mCherry(-6)-L1 | mCherry_Fwd | Same as mCherry_Fwd for GFP(-6)-H2 + mCherry(-6) |
| | | mCherry_Rev | TCATTACTTTCGAGATTTCTTG |
| | | GFP_Fwd1 | Same as GFP_Fwd1 for GFP(-6)-H2 + mCherry(-6) |
| | | GFP_Rev1 | TATCTCCTTCTTAAAGTTAATTATTTCTTCCTTCTCC TCTTTT |
| | | GFP_Fwd2 | Same as GFP_Fwd2 for GFP(-6)-H2 + mCherry(-6) |
| | | GFP_Rev2 | Same as GFP_Rev2 for GFP(-6)-H2 + mCherry(-6) |
| GFP(-6)-H2 + mCherry(-6)-L2 | GFP(-6)-H2, mCherry(-6)-L2 | mCherry_Fwd | Same as mCherry_Fwd for GFP(-6)-H2 + mCherry(-6) |
| | | mCherry_Rev | Same as mCherry_Rev for GFP(-6)-H2 + mCherry(-6)-L1 |
| | | GFP_Fwd1 | Same as GFP_Fwd1 for GFP(-6)-H2 + mCherry(-6) |
| | | GFP_Rev1 | Same as GFP_Rev1 for GFP(-6)-H2 + mCherry(-6)-L1 |
| | | GFP_Fwd2 | Same as GFP_Fwd2 for GFP(-6)-H2 + mCherry(-6) |
| | | GFP_Rev2 | Same as GFP_Rev2 for GFP(-6)-H2 + mCherry(-6) |
| GFP(-6)-H2 + mCherry(-6)-L3 | GFP(-6)-H2, mCherry(-6)-L3 | mCherry_Fwd | Same as mCherry_Fwd for GFP(-6)-H2 + mCherry(-6) |
| | | mCherry_Rev | TTATTACTTCCGCGATTTCTTCCCTC |
| | | GFP_Fwd1 | Same as GFP_Fwd1 for GFP(-6)-H2 + mCherry(-6) |
| | | GFP_Rev1 | Same as GFP_Rev1 for GFP(-6)-H2 + mCherry(-6)-L1 |
| | | GFP_Fwd2 | AGAAATCGCGGAAGTAATAACTCGAGTCTGGTAAA GAAAC |
| | | GFP_Rev2 | Same as GFP_Rev2 for GFP(-6)-H2 + mCherry(-6) |

a. This was used as the forward primer for all single-protein GFP variants.
b. All reverse primers for single-protein variants were designed to contain: 5’-XhoI cut site/stop codon/peptide/plasmid binding region-3’.
c. This was used as the forward primer for all single-protein mCherry variants.

### REFERENCES


1. Piovesan, D., Tabaro, F., Mičetić, I., Necci, M., Quaglia, F., Oldfield, C.J., Aspromonte, M.C., Davey, N.E., Davidović, R., Dosztányi, Z., et al. (2017). DisProt 7.0: a major update of the database of disordered proteins. Nucleic Acids Res *45*, D219–D227. 10.1093/nar/gkw1056.

2. Hatos, A., Hajdu-Soltész, B., Monzon, A.M., Palopoli, N., Álvarez, L., Aykac-Fas, B., Bassot, C., Benítez, G.I., Bevilacqua, M., Chasapi, A., et al. (2019). DisProt: intrinsic protein disorder annotation in 2020. Nucleic Acids Research, gkz975. 10.1093/nar/gkz975.

3. Quaglia, F., Mészáros, B., Salladini, E., Hatos, A., Pancsa, R., Chemes, L.B., Pajkos, M., Lazar, T., Peña-Díaz, S., Santos, J., et al. (2022). DisProt in 2022: improved quality and accessibility of protein intrinsic disorder annotation. Nucleic Acids Research *50*, D480–D487. 10.1093/nar/gkab1082.

4. Xue, B., Dunbrack, R.L., Williams, R.W., Dunker, A.K., and Uversky, V.N. (2010). PONDR-FIT: A meta-predictor of intrinsically disordered amino acids. Biochimica et Biophysica Acta (BBA) - Proteins and Proteomics *1804*, 996–1010. 10.1016/j.bbapap.2010.01.011.

5. Erdős, G., Pajkos, M., and Dosztányi, Z. (2021). IUPred3: prediction of protein disorder enhanced with unambiguous experimental annotation and visualization of evolutionary conservation. Nucleic Acids Research *49*, W297–W303. 10.1093/nar/gkab408.
